# MARK2/MARK3 kinases are catalytic co-dependencies of YAP/TAZ in human cancer

**DOI:** 10.1101/2024.02.26.582171

**Authors:** Olaf Klingbeil, Damianos Skopelitis, Claudia Tonelli, Aktan Alpsoy, Francesca Minicozzi, Disha Aggarwal, Suzanne Russo, Taehoon Ha, Osama E. Demerdash, David L. Spector, David A. Tuveson, Paolo Cifani, Christopher R. Vakoc

## Abstract

The Hippo signaling pathway is commonly dysregulated in human cancer, which leads to a powerful tumor dependency on the YAP/TAZ transcriptional coactivators. Here, we used paralog co-targeting CRISPR screens to identify the kinases MARK2/3 as absolute catalytic requirements for YAP/TAZ function in diverse carcinoma and sarcoma contexts. Underlying this observation is direct MARK2/3-dependent phosphorylation of NF2 and YAP/TAZ, which effectively reverses the tumor suppressive activity of the Hippo module kinases LATS1/2. To simulate targeting of MARK2/3, we adapted the CagA protein from *H. pylori* as a catalytic inhibitor of MARK2/3, which we show exerts anti-tumor activity *in vivo*. Together, these findings reveal MARK2/3 as powerful co-dependencies of YAP/TAZ in human cancer; targets that may allow for pharmacology that restores Hippo pathway-mediated tumor suppression.

## Introduction

The Hippo signaling pathway is a conserved regulator of cell identity and proliferation during metazoan development, with additional roles in tissue regeneration and in cancer progression (3). In mammals, the core of the Hippo pathway includes the kinases LATS1/2, which catalyze inhibitory phosphorylation of the YAP/TAZ transcriptional coactivators (4,5). LATS1/2 activity is, in turn, activated by MST1/2 and MAP4K kinases and by the scaffolding protein NF2, which are themselves regulated by signals from the tissue microenvironment (6–11). Once released from LATS1/2-mediated inhibition, YAP/TAZ can enter the nucleus and bind to TEAD transcription factors to activate a transcriptional program of cell proliferation and lineage plasticity (12–14).

YAP/TAZ and its upstream Hippo pathway are commonly dysregulated in human carcinomas and sarcomas to promote tumor development (1,2). This can occur via genetic (e.g. *YAP/TAZ* amplifications)(1) or non-genetic (e.g. perturbations of the extracellular matrix, metabolism, or cell polarity)(^15–19^) mechanisms, with a consequence being that many human cancers possess a powerful dependency on the function of YAP/TAZ to sustain tumor growth. Since YAP/TAZ activity is dispensable for the homeostasis of several tissues (20–22), the aberrant functioning of this pathway has motivated efforts to develop drugs that interfere with YAP/TAZ function, such as small molecules that block the interaction between YAP/TAZ and TEAD proteins (23–26). However, a major obstacle in this effort has been in identifying ‘druggable’ targets that allow for the restoration of Hippo-mediated tumor suppression in YAP/TAZ-dependent cancers.

## Results

### Paralog co-targeting CRISPR screens identify MARK2/3 as context-specific cancer dependencies

Here, we developed a dual sgRNA CRISPR vector system for performing double knockout screens of gene paralogs in search of redundant cancer cell dependencies (Fig. 1a). Using this system, we cloned a pooled library of 64,697 dual guide RNAs designed to generate 1,719 single gene knockouts and 2,529 paralog double knockouts, focusing on factors involved in signal transduction and epigenetic regulation (Fig. 1a, Supplementary Table 1,2). For each gene, we designed sgRNAs targeting exons that encode conserved protein domains to maximize the efficiency of generating loss-of-function alleles (27). We used this library to perform negative-selection screens in 22 cancer cell lines grown under standard 2D culture conditions, which represent a diverse set of tumor lineages and genotypes (Supplementary Table 3). The performance of control sgRNAs within this library supported the accuracy of these screening datasets (Supplementary Fig. S1a). For each double knockout, we quantified the degree of genetic redundancy using the GEMINI algorithm (28), which validated paralogs that are known to support cancer growth in a redundant manner, such as *HDAC1/HDAC2, ESCO1/ESCO2*, and *EP300/CREBBP* (Fig. 1b, Supplementary Table 4-6) (29–31). By excluding pan-essential paralog pairs required for all cancer cell lines tested, we nominated the kinase paralogs *MARK2* and *MARK3* as outliers showing both robust redundancy and cell line selectivity as cancer dependencies (Fig. 1b, Supplementary Fig. S1b). While prior studies have identified functions for specific MARK kinases in cancer (32–34), the essential redundant function of MARK2/3 in human cancer cells has, to our knowledge, not been previously defined.

**Fig. 1.**
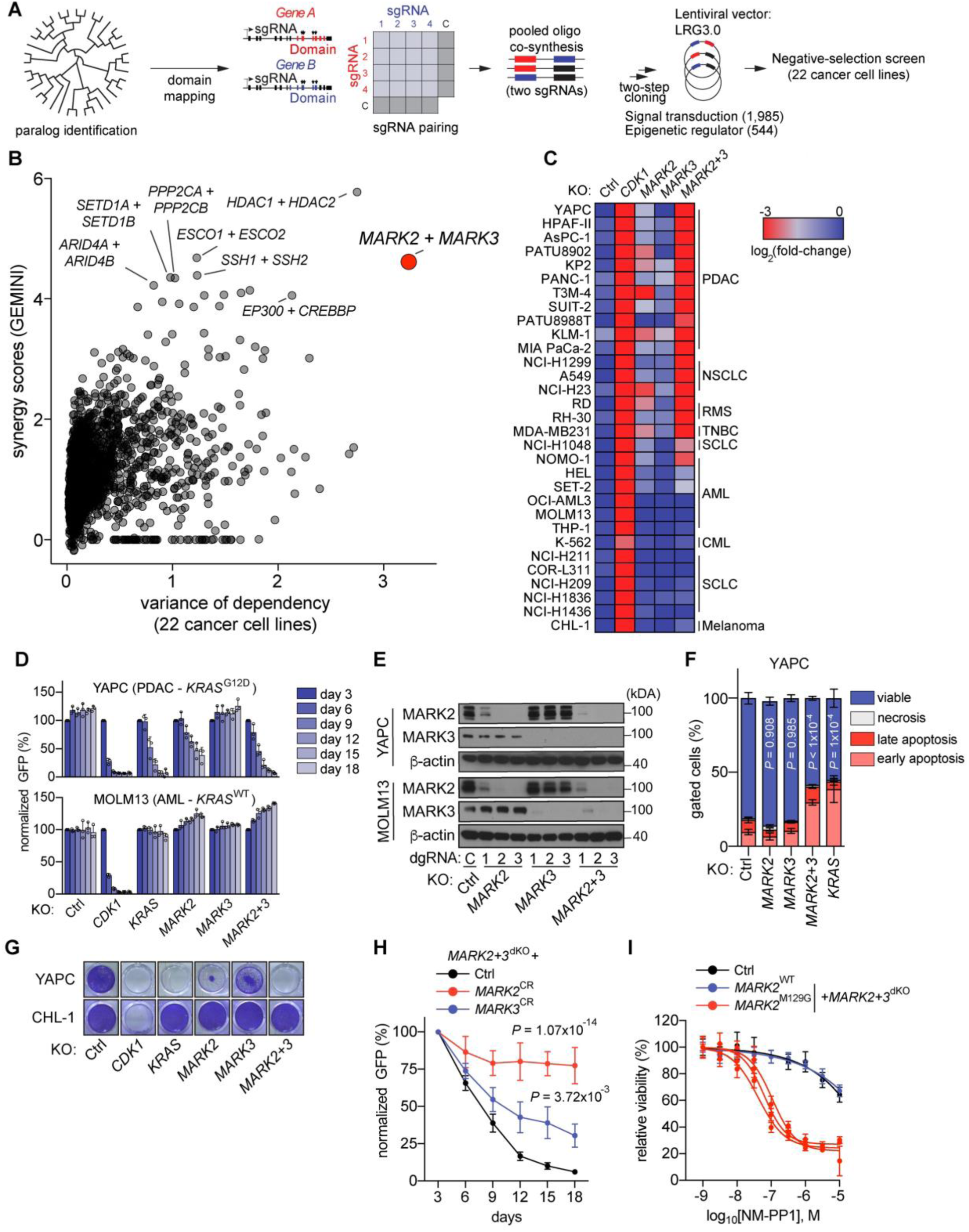
Paralog co-targeting CRISPR screens identify MARK2/3 as context-specific cancer dependencies. **A**,**B**, Workflow of paralog double knockout CRISPR screens including paralog identification, domain mapping, sgRNA design, oligo synthesis, cloning, and negative selection screening. Numbers of paralog combinations are indicated. **B,** CRISPR screening results summary, analysis of synergy between paralog gene pairs (GEMINI score) (**Supplementary Table 1,2,4-6**) maximum scores are shown together with variance of dependency (variance of average log_2_(fold-change) of double guide RNA abundance) across 22 cell lines screened. Each dot represents a double knockout paralog-pair (n=2,726) among signaling- and epigenetic regulators. **C**,**D,** Competition-based fitness assays in Cas9-expressing cancer cells after lentiviral knockout of indicated genes (expression of double guide RNAs (dgRNA) was linked to GFP). c, Heatmap color indicates the log_2_(fold-change) of normalized GFP (%GFP^+^ normalized to day 3 or 6 after infection). n=3. d, Competition-based fitness assays in the indicated cell lines. Data are shown as mean ± SD of normalized %GFP^+^ (to day 3 after infection). n=3. **E**, Western blot analysis of the indicated cell lines. **F**, Apoptosis measurements using Annexin-V and DAPI in Cas9-expressing YAPC cells. Indicated genes were knocked out using lentiviral dgRNAs linked to GFP. Data are shown as mean ± SD. n=3-6. *P* value was calculated on change in viability compared to control with one-way ANOVA and Dunnett’s correction. **G**, Crystal violet stain of indicated cells following lentiviral knockout of indicated genes. Data shown are representative of three independent biological replicates. **H**, Rescue experiment in YAPC cells using lentiviral expression of CRISPR resistant (CR) cDNAs or empty vector control (Ctrl). Data shown are the mean ± SD of %GFP^+^ (normalized to day 3 after infection). n=3. *P* values are calculated using a mixed effects model (considering the interaction of experimental groups over time) compared to Ctrl group and corrected with Bonferroni-Holm (BH). **I**, Normalized relative luminescence units (RLU) from CellTiter-Glo viability measurements of the indicated YAPC cell lines following 5 days of 1NM-PP1 treatment. Data are shown as mean ± SD. n=9 measurements from three biological replicates performed in triplicate. Four-parameter dose-response curves were plotted.

To validate these screening results, we performed arrayed-format competition-based proliferation experiments in a panel of 31 cancer cell lines (Fig. 1c, 1d, Supplementary Fig. S1c, Supplementary Table 3). These assays validated the redundancy and essentiality of MARK2/3 in 19 cancer lines, whereas 12 cancer lines proliferated normally despite effective MARK2/3 double knockout, confirmed by western blotting (Fig. 1e, Supplementary Fig. S1d). In these experiments, we noticed that MARK2/3 dependency was biased towards carcinomas and sarcomas, whereas most hematopoietic and neuroendocrine lineage cancers proliferated independently of MARK2/3 (Fig. 1c). Knockout of MARK2/3 led to a G0/G1 cell cycle arrest and apoptosis in pancreatic (YAPC) and breast (MDA-MB231) adenocarcinoma lines, with a potency that resembled the effects of inactivating the mutant *KRAS* oncogene present in these models (Fig. 1f, 1g, Supplementary Fig. S1e-g). MARK2/3 knockout in YAPC xenografts led to robust tumor growth inhibition *in vivo* (Supplementary Fig. S1h, 1i). Expression of a CRISPR-resistant *MARK2* or *MARK3* cDNA alleviated the cell fitness defect caused by the double knockout, indicating on-target effects (Fig. 1h, Supplementary Fig. S1k). Using this cDNA rescue assay, we found that mutational inactivation of kinase activity (MARK2^K82H^) compromised cancer cell proliferation (Supplementary Fig. S1j, 1l). We further validated the importance of MARK2/3 catalytic function using a bump-and-hole strategy(35), in which replacement of endogenous MARK2/3 with MARK2^M129G^, rendered the proliferation of YAPC cells sensitive to the bulky kinase inhibitor 1NM-PP1 (Fig 1i and Supplementary Fig. S1j, 1l). Collectively, these experiments validated MARK2/3 as catalytic dependencies in specific carcinoma and sarcoma cell line models.

### MARK2/3 dependency in cancer is linked to the maintenance of YAP/TAZ function

We next sought to understand why MARK2/3 is essential in some cancer contexts, but dispensable in others. Using comparative transcriptome analysis, we found that the MARK2/3 essentiality across the 31 cancer lines was highly correlated with the expression of *YAP* and *TAZ* and with the expression of canonical YAP/TAZ target genes *MYOF*, *CYR61*, *DKK1*, and *CAV1* (Fig. 2a, 2b) (36–38). Using dual sgRNA vectors, we confirmed that YAP and TAZ function redundantly as dependencies in this cell line panel in a manner that closely correlated with MARK2/3 essentiality (Fig. 2b, 2c Supplementary Fig. S2a, 2b). This observation led us to hypothesize that MARK2/3 is critical for maintaining YAP/TAZ function in diverse human cancer contexts. In support of this, we found that the inactivation of MARK2/3 led to reduced expression of a fluorescence-based TEAD:YAP/TAZ reporter in MDA-MB231 cells (Fig. 2d) (18). In addition, RNA-seq analysis performed in 20 different cancer cell line models following MARK2/3 knockout demonstrated reduced expression of a YAP/TAZ transcriptional signature in MARK2/3-dependent lines (Fig. 2e-g, Supplementary Table 7). We extended this analysis by performing genome-wide profiling of active chromatin (H3K27 acetylation), which revealed that MARK2/3 and YAP/TAZ are each critical to activate a shared set of TEAD4:YAP-bound enhancer elements (Fig. 2h, 2i, Supplementary Fig. S2c-e). Together, these results suggest that MARK2/3 are required to maintain the essential function of YAP/TAZ in human cancer.

**Fig. 2.**
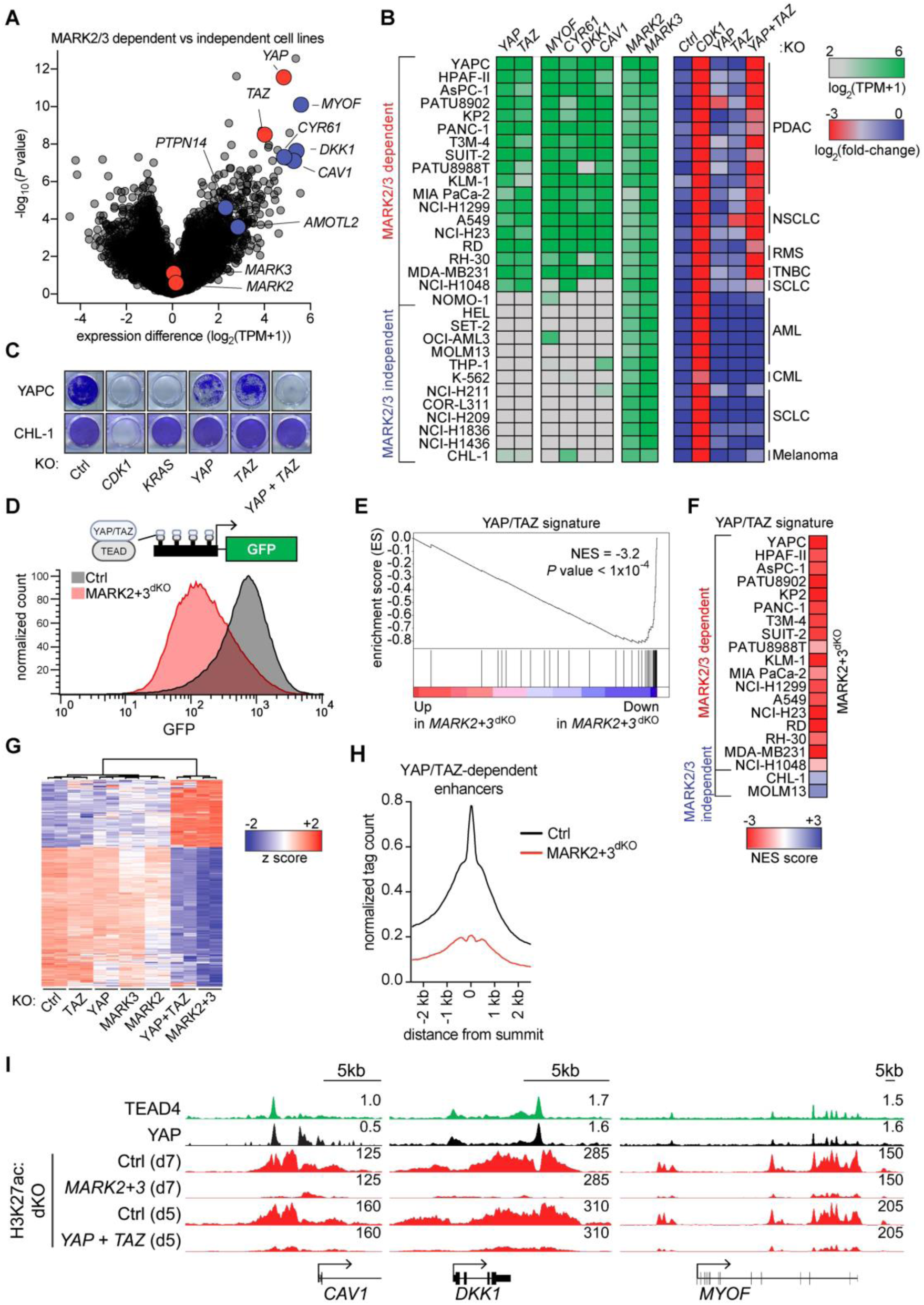
MARK2/3 dependency in cancer is linked to the maintenance of YAP/TAZ function. **A**, mRNA expression differences comparing 19 MARK2/3-dependent cell lines to 12 MARK2/3-independent human cancer cell lines. Transcriptome data were obtained from the CCLE database, KLM-1 (GSE140484) and CHL-1 (this paper). TPM, transcripts per million were calculated and the difference in log_2_(TPM+1) was plotted. *P* values were calculated using Empirical Bayes Statistics (eBayes) for differential expression with BH correction. **B**, Heatmap of MARK2/3 dependent and independent cancer cell lines showing dependence on YAP/TAZ and expression of target genes. Competition-based fitness assays in Cas9-expressing cancer cells after lentiviral knockout of indicated genes (expression of dgRNAs was linked with GFP). Heatmap color indicates the log_2_(fold-change) of %GFP^+^ (normalized to day 3 or 6 after infection). n=3. **C**, Crystal violet stain of indicated cells following lentiviral knockout of indicated genes. Data shown are representative of three independent biological replicates. **D**, Flow cytometry histogram of YAP/TAZ:TEAD reporter assay(18) in MDA-MB231 cells, on day 9 post-infection. Data are representative of three independent experiments. **E,** Gene set enrichment analysis (GSEA) of Cas9^+^ MDA-MB231 cancer cells following MARK2+3^dKO^, including normalized enrichment score (NES) and *P* value. **F,** Heatmap showing the GSEA NES for the YAP/TAZ gene signature following MARK2+3^dKO^ in dependent and independent cell lines. **G,** Heatmap of mRNA expression (log_2_(normalized count)) z-scores in Cas9^+^ MDA-MB231 cells of genes significantly down- or up regulated upon MARK2+3^dKO^. Expression values of down genes (n=188) and up genes (n=91) of two replicate samples following gene knockout were grouped based on unsupervised clustering. Significant differentially expressed genes were defined as adjusted *P* value <10^-4^ and log_2_(fold-change) >2 or <-1. *P* values from Wald test (DEseq2) adjusted using BH. **H,** CUT&RUN density profile of YAP:TEAD4 bound, YAP/TAZ^dKO^ sensitive H3K27ac marked enhancer loci (n=7,896) following MARK2+3^dKO^. Profiles shown are an average of 50bp bins around the summit of the enhancers. **i,** Occupancy profiles of public Chromatin immunoprecipitation sequencing (ChIP-seq) (TEAD4, YAP) (GSE66083) and CUT&RUN (H3K27ac) upon indicated gene knockout at YAP/TAZ target gene loci.

### MARK2/3 catalyze inhibitory phosphorylation of NF2 and activating phosphorylation of YAP/TAZ

Upon inactivating MARK2/3, we observed a striking increase in LATS1/2 T1079/T1041 phosphorylation (Fig 3a, 3b, Supplementary Fig. S3a). This activation mark is known to be catalyzed redundantly by MST1/2 and MAP4K kinases, whose activity is further enhanced by NF2 (Fig. 3a) (6). Knockout of MARK2/3 triggered reduced nuclear levels of YAP/TAZ, which is an expected outcome of strengthening LATS1/2 function (Fig. 3c). While prior studies have shown that MARK2/3 inhibits the function of MST1/2 (34,39,40), we reasoned that this substrate would be insufficient to account for the MARK2/3 dependency in cancer, since MST1/2 function redundantly with MAP4Ks to regulate YAP/TAZ in human cells (see below)(^6,8^). This prompted us to perform a broader exploration of MARK2/3 substrates in the Hippo pathway using a chemical-genetic strategy (Fig. 3d) (41). Our approach exploited gatekeeper substitutions of MARK2 (M129G) and MARK3 (M132G), which can accommodate bulky ATP-γ-S analogs (e.g. 6-Fu-ATP-γ-S). We co-expressed MARK2^M129G^ or MARK3^M132G^ with 18 different epitope-tagged Hippo pathway components in HEK293T cells, followed by treatment with 6-Fu-ATP-γ-S and immunoprecipitation-western blotting with a phospho-thio-ester-specific antibody. This approach validated the known ability of MARK2/3 to phosphorylate CDC25C and MST1/2, in accord with prior findings (Supplementary Fig. S3b-d) (34,42). In addition, we identified NF2, YAP, and, to a lesser extent, TAZ, as MARK2/3 substrates in this system (Supplementary Fig. S3b-d). Importantly, we did not detect MARK2/3-dependent phosphorylation of LATS1/2, but we detected robust phosphorylation of several MAP4K kinases (Supplementary Fig. S3b-d). To map the exact sites of phosphorylation, we performed *in vitro* kinase assays with purified MARK2 and each substrate, followed by mass spectrometric peptide quantification (Supplementary Fig. S3e-g). In these assays, MARK2 catalyzed phosphorylation on serine or threonine residues of NF2 (4 sites), YAP (5 sites), and TAZ (4 sites) (Fig. 3e-g, Supplementary Fig. S4a-k, Supplementary Table 8). By introducing alanine substitutions of these phosphosites into cDNA constructs, we confirmed the importance of these specific serine/threonine residues for MARK2-dependent phosphorylation in human cells (Supplementary Fig. S5a-e). Using mass spectrometry analysis, we also identified sites of MARK2-dependent phosphorylation on MAP4K proteins and MST1/2 (Supplementary Fig. S3g), however the known redundancy among these kinases (6) led us to prioritize NF2 and YAP/TAZ for further functional investigation (Fig. 3a).

**Fig. 3.**
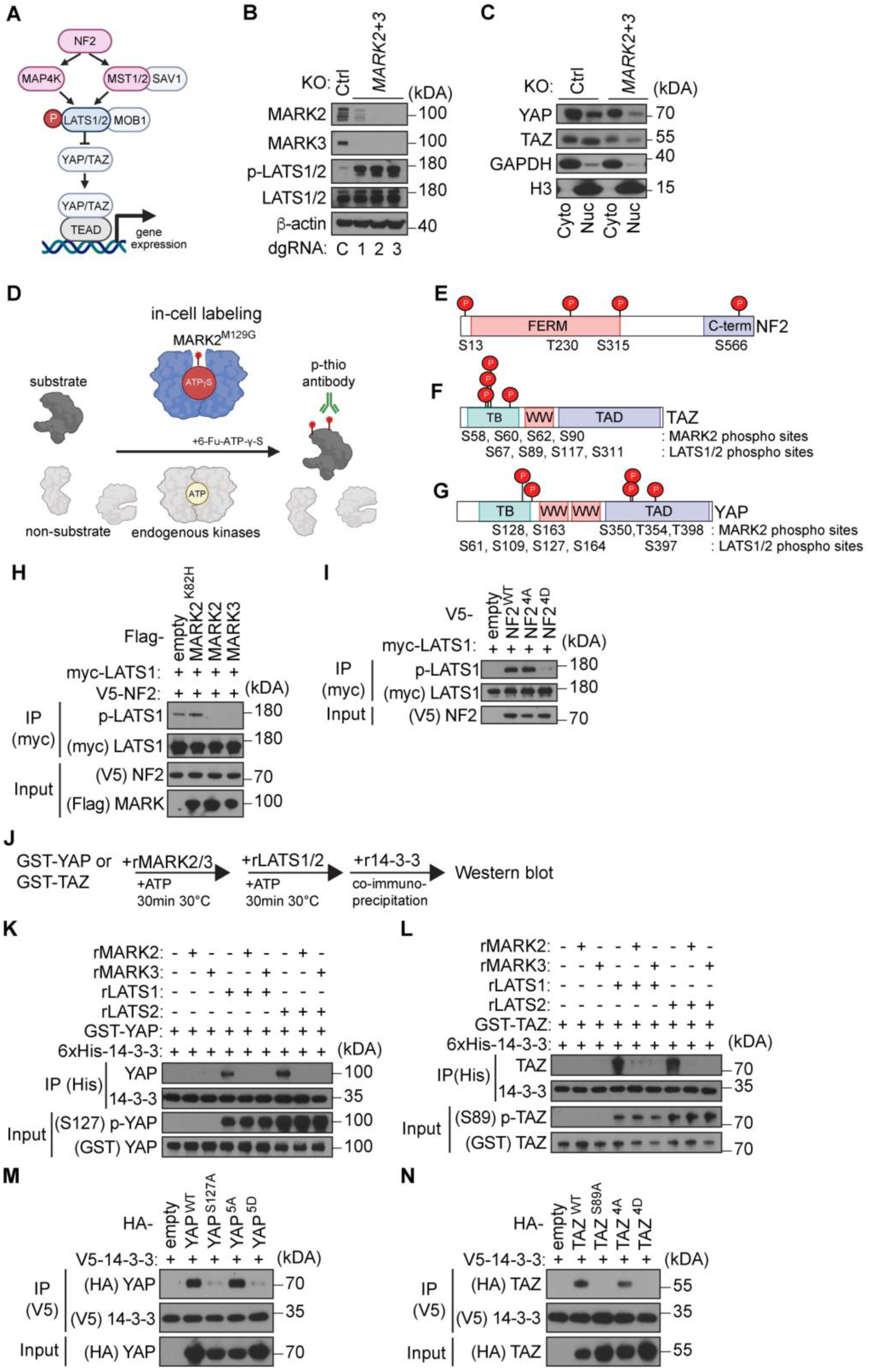
MARK2/3 catalyze inhibitory phosphorylation of NF2 and activating phosphorylation of YAP/TAZ. **A**, Illustration of the Hippo pathway. **B**,**C** Western blot analysis of Cas9^+^ YAPC cells b, whole cell lysate or c, following fractionation into nuclear (Nuc) and cytosolic (Cyto) fraction, following control^dKO^ (Ctrl) or MARK2+3^dKO^. Independent double guide RNAs (dgRNA) are indicated. **D**, Illustration of in-cell phosphorylation assay. Epitope-tagged cDNA coding for putative MARK2-substrates are transfected into HEK-293T cells together with cDNA coding for analog-sensitive mutant MARK2^M129G^. Kinase assay is performed using ATP analog (6-Fu-ATP-γ-S) selective for MARK2^M129G^. Labeled substrates are alkylated using p-nitrobenzyl mesylate (PNBM) and identified following purification by western blot analysis. **E**-**G,** Lolli-pop illustration of MARK2-dependent phosphorylation sites on NF2, YAP and TAZ identified using mass spectrometry-based phosphoproteomics. C-term=carboxy-terminal domain, TB=TEAD binding domain, TAD=transactivation domain. **H**, IP– western blot analysis evaluating the phosphorylation p-LATS1 (T1079) in presence or absence of MARK2 or MARK3 following NF2 overexpression in HEK-293T cells. Data are representative of two independent experiments. **I**, IP–western blot analysis evaluating the phosphorylation p-LATS1 (T1079) after NF2 mutant overexpression in HEK-293T cells. Data are representative of two independent experiments. **J-L,** *In vitro* phosphorylation assay and IP–western blot analysis, evaluating the interaction of 14-3-3ε and recombinant LATS1 (rLATS1) or LATS2 (rLATS2) phosphorylated GST-YAP or GST-TAZ, following phosphorylation with recombinant MARK2 (rMARK2) or MARK3 (rMARK3). Data are representative of two independent experiments. **M**, IP–western blot analysis evaluating the interaction between 14-3-3ε and YAP^5D^ (phosphomimetic mutant), YAP^5A^ (phospho-null mutant) and controls YAP^WT^ (wild type) and YAP^S127A^ (LATS1/2 phosphosite/ 14-3-3 interaction mutant) in HEK-293T cells. Data are representative of two independent experiments. **N**, IP–western blot analysis evaluating the interaction between 14-3-3ε and TAZ^4D^ (phosphomimetic mutant), TAZ^4A^ (phospho-null mutant) and controls TAZ^WT^ (wild type) and TAZ^S89A^ (LATS1/2 phosphosite/ 14-3-3 interaction mutant) in HEK-293T cells. Data are representative of two independent experiments.

Two of the sites of MARK2/3-dependent phosphorylation on NF2 were T230 and S315, which have been reported to inhibit NF2 function (43). To further evaluate this, we used a transfection-based assay in HEK293T cells (6,44), in which NF2 overexpression stimulates p-LATS1/2. We found that co-expression of wild-type MARK2/3, but not a catalytically dead mutant, negated NF2-stimulated LATS1/2 phosphorylation (Fig. 3h, Supplementary Fig. S5f). In addition, a phospho-mimetic allele of NF2, in which all four sites of MARK2-dependent phosphorylation are substituted with aspartate, was incapable of triggering LATS1/2 phosphorylation (Fig. 3i, Supplementary Fig. S5g). We also found that MARK2 was able to disrupt the physical interaction between NF2 and MAP4K kinases and block MAP4K4/6-dependent LATS1 phosphorylation (Supplementary Fig. S5h-k) (6). Knockout of MARK2/3 triggered increased levels of JUN phosphorylation, a known downstream target of MAP4K kinases (Supplementary Fig. S5l, 5m) (45). Together, our findings suggest that MARK2/3 can indirectly suppress LATS1/2 activity by directly phosphorylating upstream components of the Hippo pathway.

We next evaluated the functional importance of YAP/TAZ phosphorylation by MARK2/3. LATS1/2 have been shown to sequester YAP/TAZ in the cytoplasm by installing phosphorylation that is recognized by 14-3-3 proteins (46). Owing to the adjacent locations of several MARK2/3 and LATS1/2 substrates on YAP/TAZ (Fig. 3f, 3g) (47,48), we hypothesized that MARK2/3-dependent phosphorylation might release YAP/TAZ from 14-3-3-mediated inhibition. To evaluate this, we reconstituted LATS1/2-dependent YAP/TAZ phosphorylation using purified proteins (Fig. 3j, Supplementary Fig. S5n), which was sufficient to trigger interactions with recombinant 14-3-3ε (Fig. 3k, 3l). However, pre-incubation of recombinant YAP or TAZ with MARK2 or MARK3 and ATP eliminated the formation of 14-3-3ε complexes despite the presence of LATS1/2-dependent phosphorylation (Fig. 3k, 3l). In accord with these *in vitro* findings, expression of a phospho-mimetic allele of YAP or TAZ, in which all MARK2/3 substrates are mutated to aspartic acid, eliminated the 14-3-3ε interaction in cellular lysates (Fig. 3m, 3n). Collectively, these functional experiments support that MARK2/3-dependent phosphorylation of YAP/TAZ can disrupt the LATS1/2-dependent formation of 14-3-3 complexes.

### Regulation of NF2 and YAP accounts for the essential functions of MARK2/3 in human cancer

The biochemical findings above prompted us to perform epistasis experiments evaluating whether dual regulation of NF2 and YAP/TAZ underlies the essential function of MARK2/3 in cancer identified in our paralog screen. As expected, we found that the pharmacological inhibition or double knockout of MST1/2, or its adaptor SAV1, failed to alleviate the MARK2/3 dependency (Fig. 4a, 4b, Supplementary Fig. S6a-d). In contrast, inhibition or double knockout of LATS1/2 resulted in a bypass of MARK2/3 essentiality in four different cancer cell line models (Fig. 4a, 4c, Supplementary Fig. S6c,d). In these same models, we found that NF2 knockout or expression of a phosphomimic allele of YAP (YAP^5D^) partially alleviated the MARK2/3 dependency (Fig. 4d, 4e, Supplementary Fig. S6e). Moreover, combining the NF2^KO^/YAP^5D^ genetic alterations led to a nearly complete bypass of MARK2/3 dependency in these contexts, which resembles the effects of inactivating LATS1/2 (Fig. 4a, 4e). Collectively, these results suggest that an essential function of MARK2/3 in cancer is to regulate NF2 and YAP/TAZ, which allows for potent indirect control over the output of LATS1/2.

**Fig. 4.**
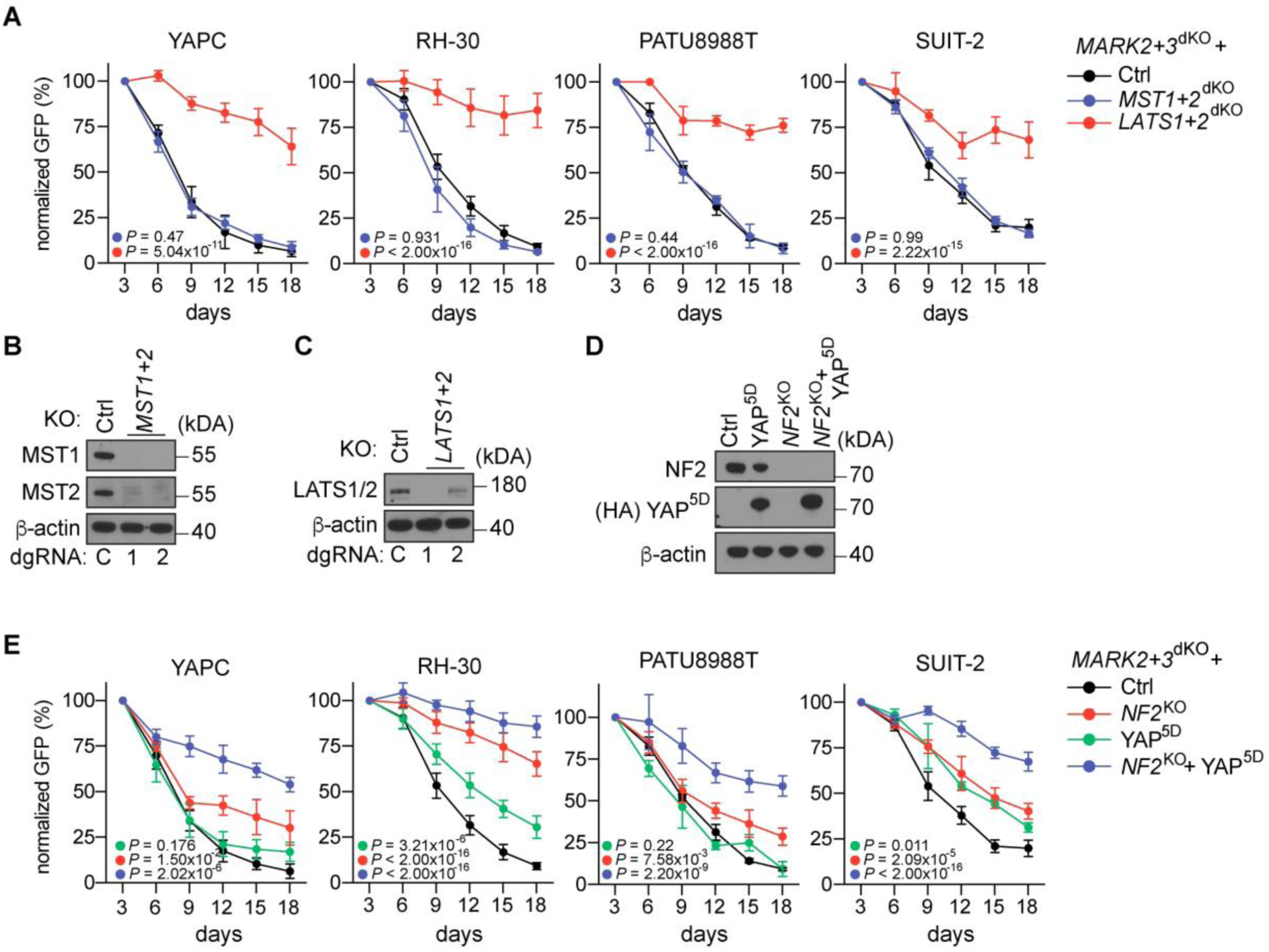
Regulation of NF2 and YAP accounts for the essential functions of MARK2/3 in human cancer. **A,** Rescue experiment of MARK2+3^dKO^ following double knockout of *LATS1*/*2* or *MST1*/*2* and control double knockout Ctrl (dgRNA targeting hROSA26 locus) in indicated Cas9^+^ cell lines. Data shown are the mean ± SD of %GFP^+^ (normalized to day 3 after infection). n=3-6. *P* values are calculated using a mixed effects model (considering the interaction of experimental groups over time) compared to Ctrl group and corrected with Bonferroni-Holm (BH). **B**,**C** Western blot analysis in YAPC cells and independent dgRNAs are indicated. **D**, Western blot analysis in Cas9^+^ YAPC cells. **E**, Rescue experiment of MARK2+3^dKO^ following knockout of *NF2* or Ctrl and lentiviral HA-YAP^5D^ overexpression. Data shown are the mean ± SD of %GFP^+^ (normalized to day 3 after infection). n=3. *P* values are calculated using a mixed effects model (considering the interaction of experimental groups over time) compared to Ctrl group and corrected with Bonferroni-Holm (BH).

### Inducible expression of a protein-based MARK2/3 inhibitor re-instates Hippo-mediated tumor suppression in organoid and xenograft tumor models

The Hippo pathway activity is known to be modulated by cell culture conditions (18), which motivated us to validate MARK2/3 dependency in tumor models with more physiological extracellular environments. Since selective small-molecule inhibitors of MARK kinases are not available, we developed a catalytic inhibitor of MARK kinase activity that could be expressed in an inducible manner in various tumor models. The EPIYA repeat region of the CagA protein of *H. pylori* was reported to potently and selectively inhibit MARK kinase activity by competing with substrate binding (49,50), a peptide we refer to here as MARK kinase inhibitor (MKI) (Fig. 5a). We observed that lentiviral expression of MKI, but not an MKI peptide harboring point mutations that abrogate MARK binding (50), reduced the nuclear levels of YAP/TAZ and suppressed the expression of a YAP/TAZ transcriptional signature (Fig 5b-e, Supplementary Fig. S7a). In addition, the proliferation arrest induced by MKI correlated with the overall sensitivity to MARK2/3 double knockout in a cell line panel (Fig. 5c). Our epistasis experiments further indicated that engineering of NF2^KO^/YAP^5D^ alleviated the sensitivity to MKI-mediated growth (Fig. 5f), thus validating MKI as a tool catalytic inhibitor that mimics the biological effects of MARK2/3 double knockout when expressed in cancer cells.

**Fig. 5.**
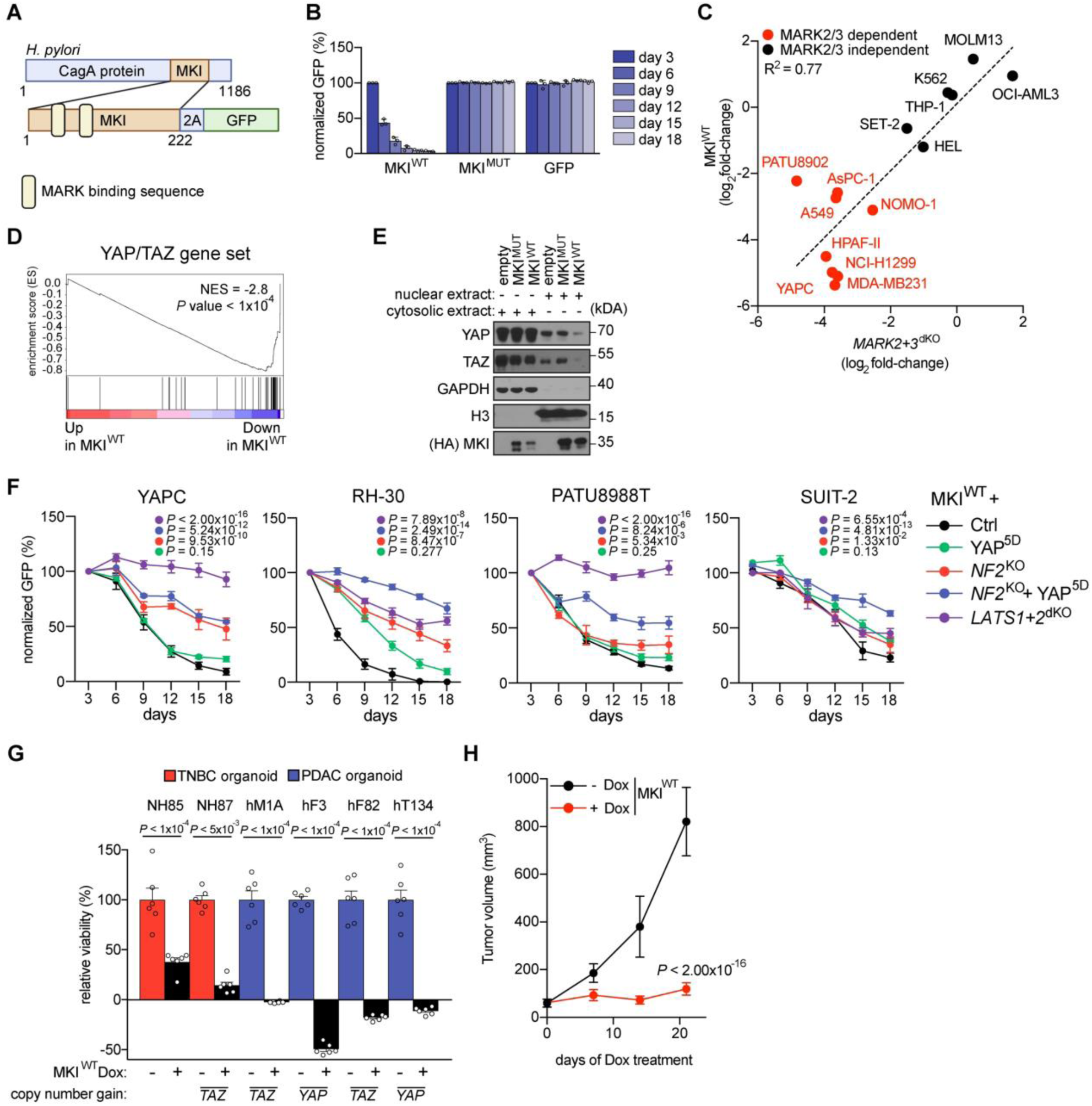
Inducible expression of a protein-based MARK2/3 inhibitor re-instates Hippo-mediated tumor suppression in organoid and xenograft tumor models. **A**, Illustration of MKI protein derived from *Helicobacter pylori* (*H.pylori*). Positioning of self-cleaving peptides (2A), GFP reporter, and number of amino acids are indicated. **B**, Competition-based fitness assays in YAPC cells after lentiviral expression of MKI^WT^ or MKI^MUT^. **C**, Comparison of log_2_(fold-change) of MKI and *MARK2+3*^dKO^double knockout competition data in Cas9^+^ cancer cell lines. Pearson correlation coefficient was calculated. Data shown are the mean of %GFP^+^ (normalized to day 3 after infection). n=3. **D**, Gene set enrichment analysis (GSEA) of RNA-seq data from MKI^WT^ compared to MKI^MUT^ expressing MDA-MB231 cells. Normalized enrichment score (NES) and *P* value are shown. **E**, Western blot analysis in YAPC cells 24h following doxycycline induced expression of indicated proteins. **F**, Rescue experiment of MKI^WT^ expression following knockout of *LATS1/2, NF2* or Ctrl (dgRNA targeting hROSA26 locus) and lentiviral HA-YAP^5D^ overexpression. Data shown are the mean ± SD of %GFP^+^ (normalized to day 3 after infection). n=3. *P* values are calculated using a mixed effects model (considering the interaction of experimental groups over time) compared to Ctrl group and corrected with Bonferroni-Holm (BH). **G**, Normalized relative luminescence units (RLU) from CellTiter-Glo viability measurements of the indicated human patient-derived triple-negative breast cancer (TNBC) or pancreatic ductal adenocarcinoma (PDAC) organoids following doxycycline (Dox) induced expression of MKI^WT^ for 10 days. Data shown are mean ± SD. n=6 measurements from two biological replicates performed in triplicate. *P* value was calculated using a two-tailed parametric t-test with Welch’s correction. **H**, Growth kinetics of subcutaneous YAPC xenografts implanted in immunodeficient mice. Expression of MKI^WT^ from a doxycycline (Dox)-inducible lentiviral construct was induced on day 10 post-injection of the cells. Data are shown as mean ± SD. n=5 per group. *P* values are calculated using a mixed effects model (considering the interaction of experimental groups over time) compared to Ctrl group (-Dox) and corrected with Bonferroni-Holm (BH).

We next engineered a vector that expresses MKI under the control of a doxycycline-inducible promoter, which was introduced into a panel of *YAP*- or *TAZ*-amplified human triple-negative breast cancer or pancreatic ductal adenocarcinoma organoid cultures. Dox-inducible expression of MKI in these models led to a strong reduction of cancer cell viability (Fig. 5g). We also introduced the dox-inducible MKI (wild-type versus mutant) expression constructs into pancreatic adenocarcinoma cells (YAPC), which were transplanted subcutaneously into immune-deficient mice. After the tumors were established (day 10), we administered doxycycline and observed that MKI, but not the point mutant control, led to a potent reduction of tumor growth *in vivo* (Fig. 5h Supplementary Fig. S7b,c). The findings validate the potent anti-tumor effects of catalytic MARK2/3 inhibition in YAP/TAZ-dependent cancers.

## Discussion

It has been observed that human cancers can be broadly classified based on the status of YAP/TAZ(51). YAP/TAZ^OFF^ tumors tend to be of hematopoietic or neural/neuroendocrine lineages, and in this context transcriptional silencing of YAP/TAZ is required for tumor development (51–53). In contrast, YAP/TAZ are activated in human carcinomas and sarcomas, which is essential for tumorigenesis (51,54). This binary classification has important clinical implications, as YAP/TAZ have powerful effects on several tumor cell phenotypes, including epigenetic plasticity and drug sensitivities (2,55). Here, we have exploited the ON vs OFF status of this pathway to reveal a strict requirement for MARK2/3 catalytic activity to support YAP/TAZ function across a diverse array of human carcinomas and sarcomas. Targeting of MARK2/3 leads to potent inhibition of YAP/TAZ and a severe compromise of tumor cell fitness; phenotypes that can be accounted for by phosphorylation of NF2 and YAP as direct MARK2/3 substrates. Our study positions MARK2/3 as dominant regulators of the human Hippo pathway, and hence a ‘druggable’ target in YAP/TAZ-dependent tumors.

Early genetic studies in model organisms implicated the MARK1-4 ortholog Par-1 as key regulator of cell polarity (56,57). Importantly, work in *Drosophila* identified Par-1 as a negative regulator of the Hippo pathway, which influences cell growth phenotypes in this organism (39). Despite this early observation, the connection between MARKs and Hippo in human cells has been controversial, with some studies suggesting MARKs can activate (34,39,40) or inhibit (33,58) YAP/TAZ function. Since these prior studies focused on the genetic manipulation of individual MARK kinase genes, genetic redundancy between MARK2/3 likely concealed the powerful inhibitory influence of human MARK kinases over the Hippo pathway. While our findings are generally consistent with the earlier *Drosophila* study(39), the mechanism by which MARK/Par-1 regulate YAP/TAZ appears to be distinct in each organism, with an expansion of upstream and downstream substrates of MARK2/3 in human cells that allow for multi-level control over the output of LATS1/2. Nevertheless, this work suggests an ancient linkage between MARK and Hippo during metazoan evolution, which may have emerged to integrate cellular polarity with organ growth and regeneration.

Prior studies have described small-molecules that block the interaction between YAP/TAZ and TEAD transcription factors (23–26,59), which are currently the most developed therapeutic strategy for targeting Hippo-dysregulated cancers (60). While the efficacy of such an approach in human patients has only recently begun to be evaluated in clinical trials (61,62), our work reveals chemical inhibition of MARK2/3 kinase activity as an alternative strategy for eliminating YAP/TAZ-addicted tumor cells. As kinases, chemical inhibition of MARK2/3 could achieve desirable selectivity and potency by leveraging decades of experience in the pharmaceutical industry at targeting this class of enzymes (63), which would differ from the challenges of modulating a protein-protein interaction (64,65). In addition, by functioning upstream to regulate LATS1/2-mediated control over YAP/TAZ, targeting of MARK2/3 would likely select for distinct resistance mechanisms from drugs targeting the TEAD:YAP/TAZ interaction (66). While the liabilities of each targeting strategy await further description in pre-clinical models and ongoing clinical studies, our study justifies consideration of MARK2/3 as an oncoprotein-like cancer target in a diverse collection of human carcinomas and sarcomas harboring hyper-active YAP/TAZ function.

## Methods

### Cell culture

The HPAF-II, AsPC-1, PANC-1, MIA PaCa-2, NCI-H1299, A549, NCI-H23, RD, MDA-MB231, NCI-H1048, NCI-H211, NCI-H209, NCI-H1836, NCI-H1436, CHL-1, OCI-AML3, THP-1, HEK-293T and K-562 were purchased from American Type Culture Collection (ATCC).

The YAPC, PATU8902, PATU8988T, NOMO-1, HEL, SET-2, RH-30, OCI-AML3 and MOLM13 cell lines were purchased from the “Deutsche Sammlung von Mikroorganismen und Zellkulturen” (DSMZ). The KP2, T3M-4, SUIT-2 and KLM-1 cell lines were purchased from the “Japanese Collection of Research Bioresources Cell Bank” (JCRB). The COR-L311 cell line was purchased from the “European Collection of Authenticated Cell Cultures” (ECACC).

All human cell lines were grown in Roswell Park Memorial Institute (RPMI) medium supplemented with 10% fetal bovine serum (FBS) and 1% penicillin/streptomycin (Gibco), if not otherwise indicated. HEK 293T and MDA-MB231 cells were grown in Dulbecco’s Modified Eagle Medium (DMEM) medium. NCI-H209, NCI-H1836, NCI-H1436, NCI-H1048 were grown in HITES medium (DMEM media supplemented with 5% FBS, 1% penicillin/streptomycin, Insulin-Transferrin-Selenium (Gibco), 10 nM Hydrocortisone (Sigma-Aldrich), 10 nM beta-estradiol (Sigma-Aldrich), 10 mM HEPES (Gibco), 2mM L-glutamine (Gibco)). All lentiviral packaging with HEK 293T cells and cancer cell line transduction was performed following standard procedures similar to those previously described (27). For organoid culture transduction, single cells were infected using a spin-infection strategy (800g for 2-4h), before virus removal and replating in Matrigel (Corning). All organoids were grown in growth factor reduced Matrigel. Human patient-derived pancreas- and breast cancer organoids were cultured in specific organoid media as described before (67,68).

### Protein lysate preparation for Western blotting and immunoblotting

Cells were lysed directly with 2x Laemmli Sample Buffer (BIO-RAD), supplemented with β-mercaptoethanol (Sigma-Aldrich) or in RIPA buffer supplemented with protease inhibitor cocktail (Roche) and Halt Phosphatase inhibitor cocktail (Thermo Fisher). The same total protein amounts or extracts from the same number of cells were loaded into each lane of an SDS-PAGE gel (NuPAGE 4– 12% Bis-Tris Protein gels, Thermo Fisher) followed by transfer to a nitrocellulose membrane. Membranes were blocked using 5% non-fat dry milk and washed using TBST following incubation both primary or secondary antibodies. After, membranes were developed with chemiluminescent HRP substrate (Pierce).

Antibodies used in this study are HRP-conjugated secondary antibodies (rabbit cytivia, NA934, 1:5,000 – 1:20,000), HRP-conjugated β-actin (Sigma-Aldrich, A3854, 1:5,000), HA (Roche, 3F10, 1:10,000), Flag (Sigma-Aldrich, A8592, 1:5,000), V5 (Invitrogen, R961-25, 1:5,000), myc (Abcam, ab62928, 1:3,000), GAPDH (Cell Signaling, D16H11, 1:3,000), H3 (Cell Signaling, D1H2, 1:5,000), GST-tag (Cell Signaling, 5475S, 1:3,000) and primary antibodies MARK2 (Abcam, ab133724, 1:1,000), MARK3 (Abcam, ab264285, 1:1,000), YAP (Cell Signaling, D8H1X, 1:1,000), p-YAP/TAZ (S127/S89) (Cell Signaling, D9W2I, 4911, 1:3,000), TAZ (Cell Signaling, E8E9G, D3I6D, 1:1,000), NF2 (Cell Signaling, D1D8, 1:1,000), MST1 (Cell Signaling, 3682T, 1:1,000), MST2 (Cell Signaling, 3952T, 1:1,000), LATS1/2 (GeneTex, GTX87014, 1:1,000), p-LATS1/2 (T1079/T1041) (Cell Signaling, D57D3, Abcam, ab305029, 1:1,000 – 1:3,000), cJUN (Cell Signaling, 60A8, 1:1,000), p-cJUN (S63) (Cell Signaling, E6I7P, 1:1,000), MOB1 (Cell Signaling, E1N9D, 1:1,000), p-MOB1 (T35) (Cell Signaling, 8699T, 1:1,000), SAV1 (Cell Signaling, D6M6X, 1:1,000), CDC25C (Cell Signaling, 5H9, 1:1,000), p-CDC25C (S216) (Cell Signaling, 63F9, 1:1,000), Thiophosphate ester (Abcam, ab92570, 1:5,000 – 1:20,000), 14-3-3 (Cell Signaling, 8312S, 1:1,000).

### Apoptosis and cell cycle analysis using flow cytometry

For Apoptosis analysis cancer cells transduced with sgRNA constructs were stained using conjugated Annexin-V proteins (Thermo Fisher Scientific) and DAPI according to manufacturer instructions. In brief, 6 days post-infection with lentivirus containing dgRNAs linked to GFP, Cells were detached and resuspended in staining buffer followed by incubation with Annexin-V and DAPI. Stained cells were analyzed by flow cytometry and data analysis was performed with FlowJo software. Early apoptotic-(Annexin-V^+^/DAPI^-^), late apoptotic- (Annexin-V^+^/DAPI^+^), necrotic- (Annexin-V^-^/DAPI^+^) and viable cells (Annexin-V^-^/DAPI^-^) were identified.

For cell cycle analysis cancer cells transduced with dgRNA constructs (day 5) were treated with 10µM EdU 4h prior to sampling. EdU incorporated into cells was stained according to manufacturer instructions (Thermo Fisher). In brief, Cells were detached and fixed in 4% PFA, permeabilized and EdU conjugated using click chemistry. Stained cells were analyzed by flow cytometry and data analysis was performed with FlowJo software. Cells were identified based on EdU signal and DNA content (DAPI).

### CRISPR screening and pooled paralog library generation

#### Library generation

The Paralog co-targeting CRISPR library was optimized for the use of SpCas9, a system we recently published(69). Oligonucleotide pools (n=64,697) double guide RNAs targeting 1,719 single gene and 2,529 gene combinations were synthesized (Twist Bioscience) with BsmBI cutting sites in between overhang sequences for the dual crRNA fragment. Primers matching the overhang for the lentiviral backbone were used to amplify the oligonucleotide pools. PCR products were purified and cloned using Gibson assembly master mix (New England BioLabs) into LRG3.0, a lentiviral vector with human U6 and bovine U6 promoters expressing the two sgRNAs in inverse orientation. To incorporate the dual tracrRNA, the purified tracrRNA fragment was cloned in between the dual crRNAs by a second round of Gibson assembly.

#### Paralog library screening

To generate stable cell lines, cells were first transduced with a Cas9 vector (Addgene: 108100). Next, cell lines were transduced with the paralog co-targeting CRISPR library virus aiming for a representation of 1,000 cells per sgRNA at a low multiplicity of infection (MOI ∼0.3). Briefly, cell lines were transduced by spin infection for 45 min at 600g. On day 3, an initial sample was taken and cells were re-plated maintaining representation. Once 10 cell doublings were reached samples for genomic DNA extraction were again taken.

#### Genomic DNA extraction

Cells lysed in extraction buffer (10mM Tris, 150mM NaCl, 10mM EDTA, Proteinase K (0.02mg/mL), SDS (0.1%)). Lysates were incubated at 56°C for 48h and genomic DNA was extracted using TRIS-saturated phenol (Thermo Fisher Scientific).

#### dgRNA PCR for Illumina sequencing

DNA was PCR-amplified and barcoded with P5/P7 primers (Integrated DNA Technologies) using Taq-Gold DNA polymerase (Thermofisher) according to the manufacturer’s instructions. Briefly, Taq polymerase, reaction buffer, Magnesium chloride, primers, and 1µg of genomic DNA were mixed and used for each reaction (round 1: PCR for 11 cycles). Amplified DNA was size selected (200-300bp) and barcoded in a second round PCR using stacked P5/P7 primers (round 2: PCR for 9 cycles). The PCR product was sequenced using a paired-end 75 base pair (bp) reads protocol (Illumina).

#### Calculation of paralog CRISPR screening log_2_(fold-change), synergy, P value and FDR

Reads were counted by mapping dgRNA sequences to the reference file of the library and a pseudo count of 16 was added. The GEMINI R (v.1.4.0) package was used to calculate log_2_(fold-changes) (LFC) and synergy scores and statistics with their corresponding P and FDR values (Supplementary Table 2,4-6). In brief, GEMINI calculates the LFC of the dgRNA abundance between initial time point (average abundance of dgRNAs day3 n=10) and the 10-doubling time endpoint. GEMINI has been used to compute the synergy score by comparing the LFCs of each gene pair to the most lethal individual gene of the pair. Non-synergistic pairs were used to calculate FDR and P value. Bayesian analysis and the prior choice were performed as described previously (28).

#### Paralog gene identification and functional domain mapping

Paralog pairs were identified by aligning human proteome (>100,000 amino acid sequences) using the Basic Local Alignment Search Tool (BLAST). Matches originating from the same gene were removed. Each top-scored paralog-pair identified (*E* value < 0.01), that shared the same functional domain of interest was included in the Paralog library. In addition, high-scoring paralogs (*E* value < 10^-100^) were included. Functional domains were mapped using Reverse Position-Specific BLAST and the conserved domain database (CDD) (70).

#### Selection of sgRNAs and controls

Domain annotation and sgRNA positions were compared and sgRNAs cutting in functional domain regions were included in the sgRNA selection pool. sgRNAs with off-targets in paralog genes were removed from the selection pool. Additionally, sgRNAs incompatible with the cloning strategy were removed from the selection pool. sgRNAs were picked based on their off-target score (calculated based on the number of off-target locations in the human genome factored by the fall-off in cutting-efficiency of spCas9 in case of crRNA sequence miss-match). For each gene, 3-4 selective domain-focused sgRNA were picked and combined. A set of sgRNAs targeting known essential genes as positive controls (n=28) and a set of non-targeting (n=97) as well as non-coding region targeting negative controls (n=54) were included in the library. To construct cell line-specific negative controls (non-synergistic pairs), we selected genes that were not expressed in a cell line according to the RNA-seq data (log2(TPM + 1) < 0.1).

### Arrayed GFP competition assays

For validation, two sgRNAs were synthesized together with bovine U6 promoter as gene blocks (Integrated DNA Technologies) and cloned using Gibson assembly into LRG2.1T (Addgene, 65656). All inserts were verified by Sanger sequencing (Eurofins Genomics). To generate LATS1/2 and MST1/2 double knockout pools two sgRNAs co-targeting LATS1/2 or MST1/2 were combined and two sgRNA targeting SAV1, NF2 were combined on one vector. For lentivirus packaging, HEK 293T cells were transfected with sgRNA, pVSVg, psPAX2 plasmids (Addgene, 12260) using PEI reagent (PEI 25000). Percent GFP^+^ populations were followed over time after infection using the Guava Easycyte flow HT instrument (Millipore). Complete sgRNA sequences are given in Supplementary Table 9.

### Generation of ectopic overexpression vectors

All cDNAs were either cloned from Addgene plasmids or synthesized as indicated below. CRISPR-resistant cDNAs were generated either by mutating the PAM sequence or sgRNA binding sites into synonymous codons. All cDNAs were cloned into lentiviral constructs derived from LentiV (Addgene 108100), altered to contain internal ribosome entry site (IRES) elements and selection marker resistance genes. For doxycycline induction of cDNA expression, genes were cloned into Doxi-LentiV (derived from Addgene, 80921, 89180 and 71782) vectors and expression was induced using 2 µg/ml doxycycline.

MARK2 (Addgene, 23404) and MARK3 (Addgene, 23716) were cloned into the LentiV-IRES vector after the addition of a Flag tag at the N terminus. Hippo pathway genes-LATS1, LATS2, NF2, SAV1, TAZ, MOB1A, MOB1B, MST1, MST2, TEAD1, YAP and GFP, CDC25C, YWHAE (14-3-3ε) encoding V5, HA or myc-tagged cDNAs were from Addgene (66851, 66852, 32834, 32836, 32839) or synthesized (IDT). cDNA encoding for MAP4K1, MAPK4K2, MAPK4K3, MAPK4K4, MAPK4K5, and MAPK4K6, were from Addgene (23484, 23644, 23664, 23486, 23611, 23522) 3xHA tagged and cloned into LentiV. The MAPK4K7 expression vector was built by Vector Builder. All mutations were introduced by geneBlock synthesis or PCR. MKI^WT^ was derived from the coding sequence of CagA (*H.pylori* strain 26695). The sequence containing the EPIYA-repeat regions amino acid position 885-1105 was codon optimized. The cDNA was synthesized and cloned into LentiVi-P2A-GFP or Doxi-LentiV after the addition of a 3xHA or Flag tag at the N terminus. To generate a mutant of MKI with impaired MARK binding capacity (MKI^MUT^) the leucine 109/143 in the two MARK binding motifs of MKI^WT^ were mutated to glycine.

### Generation of TEAD binding reporter linked to GFP

To generate a TEAD-driven GFP reporter, the promoter of the established TEAD binding reporter (8xGTIIC)(18) (Addgene, 34615) was fused into a construct containing destabilized GFP (Addgene, 138152).

### Generation of clonal analog sensitive YAPC cells for growth assays

MARK2 analog-sensitive mutants were generated by mutating the gatekeeper amino acid methionine 129 to glycine. The functionality of this mutant was confirmed using rescue assays. YAPC cells were infected with cDNA CRISPR resistant to sgMARK2+3 and 3 single cell clones were picked. Mutation of endogenous MARK2 and MARK3 locus for all clones was confirmed using genotyping methods (PCR and nanopore sequencing).

### Cloning, expression, and purification of recombinant proteins

ORF encoding human MARK2 (Addgene, 23404) was cloned into pFL system with an N-terminal Strep2SUMO tag. Bacmid was generated using pFL vector using DH10MultiBac cells (Geneva Biotech). Sf9 cells were transfected with purified bacmids. Cells were lysed and rMARK2 was purified using StrepTactin Super flow resin. Protein was aliquoted and snap-frozen at −80°C. Protein concentration was estimated by measuring Abs_280nm_ and samples were assessed by Coomassie staining and MS analysis, confirming the absence of other protein kinases. Recombinant LATS1, LATS2, MARK3 and 14-3-3ε were purchased (Active Motif, 81209, Signalchem, L02-11G, M45-10G, Y75-30H) and purity, correct protein size was confirmed by Coomassie staining.

Human ORFs encoding YAP and TAZ were cloned into pGEX4T1 vector with N-terminal GST-tag. BL21-CodonPlus (DE3)-RIPL competent cells (Agilent, 230280) are transformed with sequence-validated vectors. Protein expression was induced with IPTG (GoldBio, I2481C) at 16°C for 18 hours. Bacteria were sedimented, lysed, sonicated and cleared lysates were loaded, washed followed by elution using (50 mM Tris pH 8, 300 mM NaCl, 10% glycerol, 20 mM reduced L-glutathione). Purified proteins were aliquoted and flash-frozen at −80°C. The purity of the proteins was assessed by Coomassie staining. Protein concentration was estimated through Abs_280nm_ measurements.

### In-cell phosphosubstrate identification

Gatekeeper mutant MARK2^M129G^ or MARK3^M132G^ cDNA was co-transfected together with cDNAs of individual genes into HEK 293T using polyethyleneimine (PEI). After 24h cells were harvested and incubated for 30 min at 30°C in bulky-ATP-analog (N⁶-Furfuryl-ATP-γ-S) containing Kinase-labeling buffer (Protease inhibitor, 20 mM HEPES, 100 mM potassium acetate, 5mM sodium acetate, 2mM magnesium acetate, 10 mM magnesium chloride, 1mM EGTA, 45 µg/mL Digitonin, 0.5 mM TCEP, 5mM GTP, 600 µM ATP, 75 µM N⁶- Furfuryl-ATP-γ-S). Cells were lysed using RIPA buffer (with the addition of 0.1% SDS and 250 U/mL Benzonase). Thiophosphorylated substrates were alkylated using 2.5 mM para-nitrobenzyl mesylate (PNBM) for 10min at RT. Target proteins were affinity purified and analyzed using western blot and anti-thiophosphate ester-specific antibodies.

### Identification of phosphosites using mass spectrometry (MS) and phosphoproteomics

#### Sample preparation and MS recording

Substrate cDNAs were transfected into HEK 293T as described above and sampled 24 h after transfection. Samples were affinity purified using HA-agarose beads (Sigma-Aldrich) and treated with 800 U of Lambda Phosphates (New England Biolabs) for 30 min at 30°C. Beads were washed with RIPA buffer (with Protease inhibitor and Phosphatase inhibitor cocktails). Next, beads bound proteins were incubated for 30min at 30 °C with 3 µg rMARK2 in Kinase-buffer (Tris-HCl pH=7.5, 5 mM MgCl2, 2 mM EGTA, 0.5 mM DTT, 100 µM ATP, Protease- and Phosphatase inhibitor cocktail). Phosphorylated substrates and negative controls were resolved by SDS-PAGE and proteins were stained with Coomassie blue. The bands corresponding to each putative substrate were excised, and gel bands were de-stained. After irreversible alkylation of Cysteine residues, proteins were digested with Trypsin, and peptides were analyzed by LC-MS/MS. Peptides were resolved by nanoscale reversed-phase chromatography and ionized by electrospray (2,200V) into a quadrupole-orbitrap mass spectrometer (Thermo Exploris 480). The MS was set to collect 120,000 resolution precursor scans before data-dependent HCD fragmentation and collection of MS/MS spectra. The area under the curve for chromatographic peaks of precursor peptide ions was used as quantitative metrics for label-free quantification.

#### Identification of phosphosites

Raw files were analyzed using the Proteome Discoverer environment. For peptide identification, spectra were matched against the UniProt human sequence database, supplemented with common contaminants from the cRAP database and with the sequences of the recombinant proteins expressed as substrates. S/T/Y phosphorylation, N/Q deamidation, and M oxidations were set as variable modifications. Alkylation of C residues with CEMTS was set a static modification. Up to 3 missed trypsin cleavages were allowed. Peptide-spectral matches were filtered using Percolator to maintain 1% FDR using the target-decoy method. The area under the curve defined by peptide ion XIC was integrated and used as a quantitative metric for label-free quantification. To evaluate differential phosphorylation in MARK2-treated samples compared to controls, peptides from each putative substrate were parsed out, and label-free quantification (LFQ) AUC values were used as metrics for relative chemical isoform abundance across conditions. Peptides with no LFQ value in any of the samples were disregarded. For peptides only quantified in one experimental arm, the missing value was imputed using a value smaller than the smallest empirical LFQ in the dataset (value chosen as a proxy for LFQ at detection limit). Relative amounts of phosphorylated peptides in MARK2 treated and control samples were assessed for each chemical isoform independently. Phosphopeptides that were either specifically detected in the MARK2 treated samples or showing differential abundance across conditions (>2-fold-change in MARK2 treated vs untreated sample) and whose identity could be confirmed by manual spectral interpretation were prioritized for further validation using in-cell phosphosubstrate identification strategy described above. The fragmentation spectra supporting peptide identity and phosphorylation localization together with the extracted precursor ion chromatogram (XIC) can be found in Supplementary material MS.

### Crystal violet staining

Cas9-expressing cancer cells were infected with lentivirus. After 3 days GFP percentage was determined using flow cytometry. GFP^+^ cells were seeded into 24 well plates at a density of 5,000/well. Cells were selected and grown for 10-12 days in the presence of 10µg/mL Blasticidin for controls to reach near confluency. Media was changed every 3 days. Cells were fixed using 4% paraformaldehyde for 15 min followed by staining with Crystal violet (1mg/mL in 90/10% Water/Ethanol) for 5 min. Wells were washed 4 times with water and plates were imaged.

### Subcellular fractionation assay

Following perturbation, cancer cells were treated with 500µM cytosolic extraction buffer (10mM HEPES, 10mM KCl, 1mM DTT, 0.1 mM EDTA, 0.1mM EGTA) for 10min on ice. Cells were vortexed for 10sec after the addition of NP40 (final 0.65%) to allow hypotonic cell membrane lysis, followed by 5 min 1,500 g centrifugation at 4°C. Cytosolic fraction was removed and pelleted nuclei were lysed in RIPA buffer supplemented with 250 U/mL Benzonase and Protease- and Phosphatase inhibitor cocktail.

### Co-Immunoprecipitation assays

HEK 293T cells were transfected with vectors expressing myc-LATS1, myc-LATS2, V5-14-3-3 or V5-NF2 together with Flag-MARK2, Flag-MARK2^K82H^ or Flag-MARK3 and wild-type or mutant HA-tagged substrate cDNAs. For immunoprecipitation, cells were lysed in NP40 buffer (20 mM of Tris-HCl, 100 mM of NaCl, 1% NP40, 2 mM of EDTA, Protease- and Phosphatase inhibitor cocktail) or RIPA buffer (Thermo Fisher Scientific) for 10 min at 4 °C. Protein lysates were then centrifuged at 13,000g for 15 min at 4 °C. The supernatant was then transferred to new collection tubes and incubated with to 30 µl of prewashed anti-myc or -V5 beads (Chromotek) and equilibrated to a final volume of 1000 µl by adding lysis buffer. Precipitation was performed at 4 °C overnight and washed 4-5 times with lysis buffer. Samples were eluted by boiling for 10 min in 2x Laemmli Sample Buffer supplemented with β-mercaptoethanol.

### In vitro phosphorylation and interaction assay

Bacterial purified recombinant GST-YAP or GST-TAZ were pre-incubated for 30min at 30 °C with recombinant MARK2 or MARK3 in Kinase buffer followed by incubation with either recombinant LATS1 or LATS2 for an additional 30min. Phosphorylated YAP or TAZ were then incubated with 6xHis-14-3-3 bound to Ni-NTA affinity resin for 4-16h followed by washing and samples elution.

### RNA-seq, CUT&RUN sample preparation and library construction

For RNA-Seq libraries, total RNA was prepared using TRIzol reagent according to the manufacturer’s protocol (Thermo Fisher Scientific). Libraries were constructed with the TruSeq Sample Prep Kit v2 (Illumina) following the manufacturer’s protocol. Briefly, 2 μg of total RNA was used for Poly-A enrichment, fragmentation, cDNA synthesis, end repairing, A tailing, adapter ligation and library amplification. For CUT&RUN, antibody-guided DNA cleavage was performed using the CUTANA CUT&RUN kit (EpiCyper) according to the manufacturer’s instructions. Briefly, 500,000 knockout cells were crosslinked for 1 min using 1% paraformaldehyde (PFA) and quenched using Glycine for min. Pre-washing buffer was used with detergents (0.05% SDS and 0.2% Triton X-100). Antibodies used were H3K27ac and IgG (EpiCyper, 13-0045;13-0042). Libraries were constructed with the NEBNext Ultra II DNA Library Prep Kit (New England BioLabs) following the manufacturer’s low DNA protocol. Briefly, complete CUT&RUN DNA extracts were spiked-in with E. coli DNA fragments and subjected to end repair, A tailing and adapter ligation (at 1/25 dilution) followed by PCR amplification. Libraries were purified using AMPureXP beads before and after PCR. Barcoded libraries were sequenced using an Illumina Nextseq.

### Bioinformatics- RNA-seq, GSEA, ChIP-seq analysis

#### Basal expression levels, copy number variations and mutations

For cell lines basal expression data (TPM) and copy number variations (CNV) absolute values from the cancer cell line encyclopedia (CCLE)(71) were used. RNA-seq data for KLM-1 was obtained from GSE140484. Mutational information from both the CCLE and Cosmic databases was used (72). TNBC and PDAC organoid CNV data were previously published (67)(68).

### RNA-Seq analysis

Raw reads were pseudo-aligned to the transcriptome of the human genome (hg38) using Kallisto (73) with bootstrap 100. For differential gene expression analysis, pseudoalignment counts were read into DESeq2, comparing samples vs control (Ctrl^KO^) with two replicates for each sample. The differential expression gene analysis was performed using a gene expression cutoff of >0.5 TPM. Results from multiple sequencing runs were batch-corrected using the R package (sva), before count normalization, transformation, and z-score calculation. For heatmap, z-scores of normalized counts from significantly (adjusted P value < 10^-4^) down or up-regulated (log_2_(fold-change) < −1 or > 2) genes in MARK2+3^dKO^ condition were used and plotted using R package (ComplexHeatmap).

### CUT&RUN and ChIP-seq analysis

Raw reads were aligned to the human genome (hg19) and e.coli genome (K12) using Bowtie2 software in sensitive mode(74). Duplicate reads were removed before peak calling. Deeptools was used to normalize samples to e.coli-DNA spike-in controls. Peaks were identified using MACS2 software (75) using 5% FDR cut-off and broad peak option for histone or narrow peak option for transcription factor-ChIP-seq datasets. H3K27ac peaks identified from Ctrl^KO^ and MARK2+3^dKO^, YAP+TAZ^dKO^ samples were merged and overlapping peaks were combined. Normalized tag counts were calculated using the Bamliquidator package (https://github. com/BradnerLab/pipeline) without read extension and log_2_(fold-change) between control and dKO samples was calculated for each peak. YAP/TAZ sensitive enhancers were defined by bound by H3K27ac signal reduction (−1.5 > log_2_(fold-change)) and binding of YAP and TEAD4 in ChIP-seq (only enhancers with relative tag count >3 in Ctrl samples were used; n=7,896; Supplementary Table 11).

ChIP-seq datasets of TEAD4 and YAP from MDAMB231 cells were obtained from public GEO data sets TEAD4 and YAP (GSE66081). Sequencing depth normalized ChIP-seq and CUT&RUN pileup tracks were generated using the UCSC genome browser.

### Generation of YAP/TAZ gene signature and gene set enrichment analysis (GSEA)

The differential gene expression gene lists of YAP+TAZ^dKO^ compared to Ctrl^KO^ were ranked and the top 200 downregulated genes in YAP+TAZ^dKO^ condition were combined. Gene counts were ranked and genes found in at least 1/3 of models were used to generate a general cancer cell line YAP/TAZ target gene set (n=43) (Supplementary Table 7). Differentially expressed gene lists were further analyzed using gene set enrichment analysis with a weighted GSEA Pre-ranked tool. 1,000 gene set permutations were applied(76) and the common cancer YAP/TAZ target gene set was used to analyze the effects of sgMARK2/3 double guide RNAs on gene expression. All fold-changes are provided in Supplementary Table 10.

### *In vivo* tumor growth assay

For tumor growth models, cells were injected into the left or right flank. For Dox-inducible MKI cDNA transduced cells mice were. For conditional MARK inhibition experiments in vivo, 1×10^5^ TRE3G-MKI^WT/MUT^-PGK-rtTA3 cancer cells in 100µL growth factor reduced Matrigel were transplanted subcutaneously into the left or right flank of NOD.Cg-Prkdc^scid^ Il2rg^tm1Wjl^/SzJ (NSG) mice. Animals were treated with doxycycline in either drinking water (2 mg/ml with 2% sucrose; Sigma-Aldrich) to induce MKI protein expression. For stable knockout experiments *in vivo*, YAPC cells were transduced with (hU6-sgRNA-bU6-sgRNA)-EFS-GFP-2A-BlastR lentivirus, followed by selection with Blasticidin for 3 days. After, 1×10^5^ GFP^+^ viable cells were transplanted subcutaneously in 100µL growth factor reduced Matrigel into the right flank of NSG mice. For all subcutaneous xenograft experiments tumor growth was monitored using caliper measurements. The humane study end-point was determined as the control group’s average tumor size reaching > 600 mm^3^.

### Proliferation, viability assay

For the proliferation assays, cells were seeded at a density of 500 cells per well into 96-well plates. Cells were treated 24h after seeding and cell viability was assessed 5 days after treatment using the Cell Titer-Glo luminescent cell viability assay (Promega). Cells treated with vehicle control DMSO (0.1%) or killing control 10µM proteasome inhibitor (MG132). Percent viability was calculated by normalizing RLU to DMSO (0.1%) after subtraction of killing control MG132 (10µM) signal.

For organoids, 5,000 or 10,000 cells were seeded in a 10% Matrigel/90% organoid media mix and grown for 10 days in the presence or absence of 2µg/mL doxycycline, before assessment of viability using the Cell Titer-Glo luminescent cell viability assay (Promega).

### Animal studies

All mouse experiments were approved by the Cold Spring Harbor Animal Care and Use Committee. Animals were treated with doxycycline in drinking water (2 mg/ml with 1% sucrose; Sigma-Aldrich) to induce cDNA expression.

---

**Supplementary Fig. S1.**
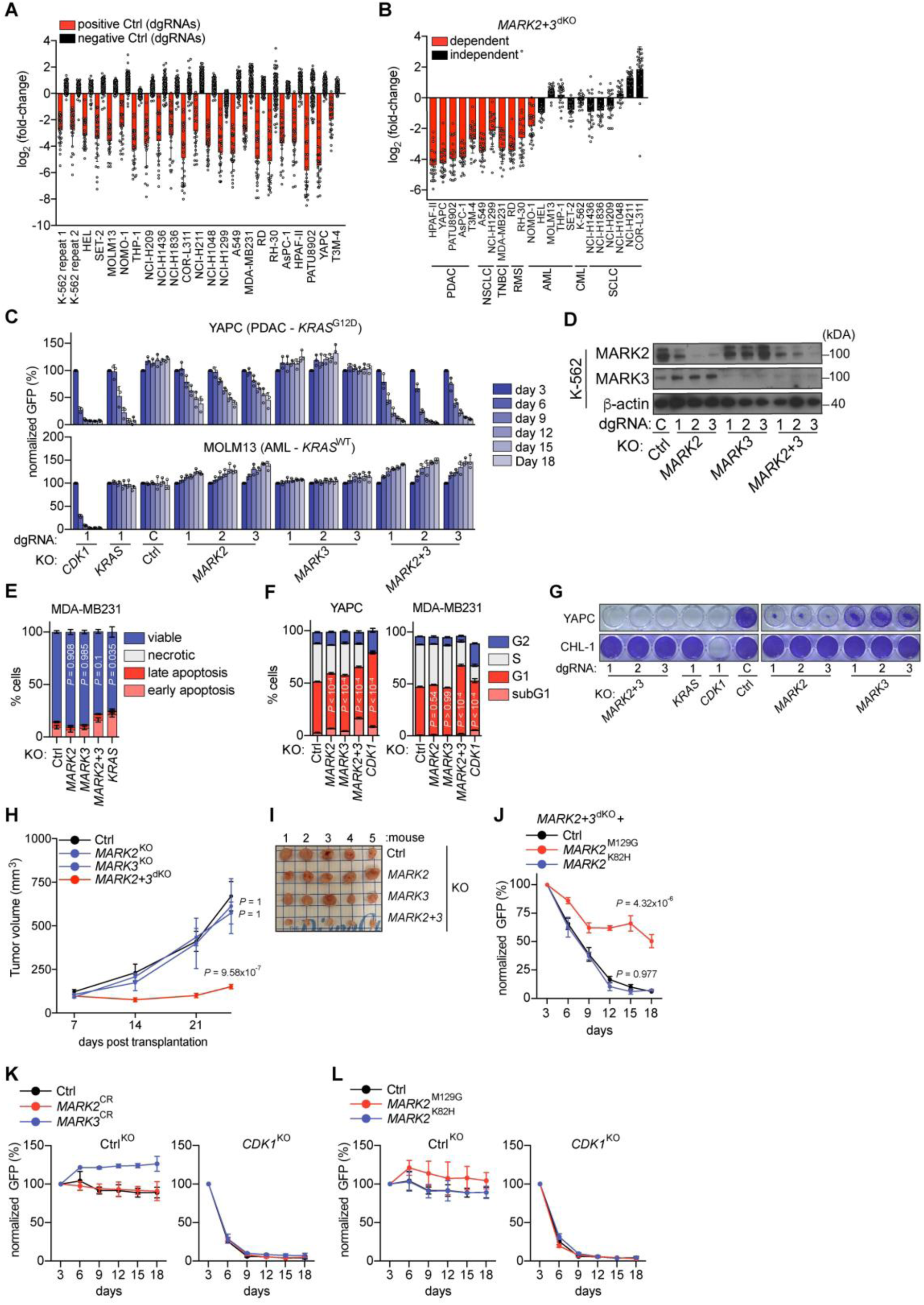
**A,B** CRISPR screening results from 22 cancer cell lines. **A**, Abundance fold-change of positive controls (dgRNAs targeting essential genes n=28 paired with control) and negative controls (dgRNAs targeting non-coding regions n=54 and nontargeting dgRNAs n=97). Data are shown as mean ± SD **B,** Abundance fold-change of dgRNAs targeting MARK2+3. Each dot represents a single dgRNA. Data are shown as mean ± SD n=24 dgRNAs. **C,** Competition-based fitness assays in Cas9-expressing cancer cells after lentiviral knockout of indicated genes with independent dgRNAs (expression of dgRNAs was linked with GFP) (Data shown are an extension of Fig. 1D). Data shown are the mean ± SD of %GFP^+^ (normalized to day 3 after infection). n=3. **D**, Western blot analysis of Cas9^+^ K-562 cells. **E**, Analysis of apoptosis assay using Annexin-V and DAPI in Cas9^+^ MDA-MB231 cells. Indicated genes were knocked out using lentiviral dgRNAs linked to GFP. Data are shown as mean ± SD. n=3-6. *P* value was calculated on change in viability compared to control with one-way ANOVA and Dunnett’s correction. **F,** EdU incorporation assays following indicated gene knock out using lentiviral dgRNAs linked to GFP in Cas9^+^ indicated cells. Data are shown as mean ± SD. n=3. *P* value was calculated on change in S-phase population to control with one-way ANOVA and Dunnett’s correction. **G**, Crystal violet stain of indicated cells following lentiviral knockout of indicated genes. Data shown are representative of three independent experiments and an extension of Fig. 1G. **H**, Growth kinetics of subcutaneous YAPC xenografts implanted in immunodeficient mice. Indicated genes were knocked out just before injection. Data are shown as mean ± s.e.m. n=5 per group. *P* values are calculated using a mixed effects model (considering the interaction of experimental groups over time) compared to Ctrl group and corrected with Bonferroni-Holm (BH). **I,** Tumor imaging at the end-point of the xenograft experiments shown in **H**. **J**, Rescue experiment in Cas9^+^ YAPC cells using lentiviral overexpression cDNA of CRISPR resistant (CR) analog sensitive mutant *MARK2^M129G^*, kinase-dead mutant *MARK2^K82H^* or empty vector control (Ctrl). Data shown are the mean ± SD of %GFP^+^ (normalized to day 3 after infection). n=3. *P* values are calculated using a mixed effects model (considering the interaction of experimental groups over time) compared to Ctrl group and corrected with Bonferroni-Holm (BH). **K**,**L** Competition-based fitness assays for Ctrl (dgRNA targeting hROSA26 locus) and knockout of essential gene *CDK1* corresponding to experiments shown in Fig. 1H and **J**. Data shown are the mean ± SD of %GFP^+^ (normalized to day 3 after infection). n=3.

**Supplementary Fig. S2.**
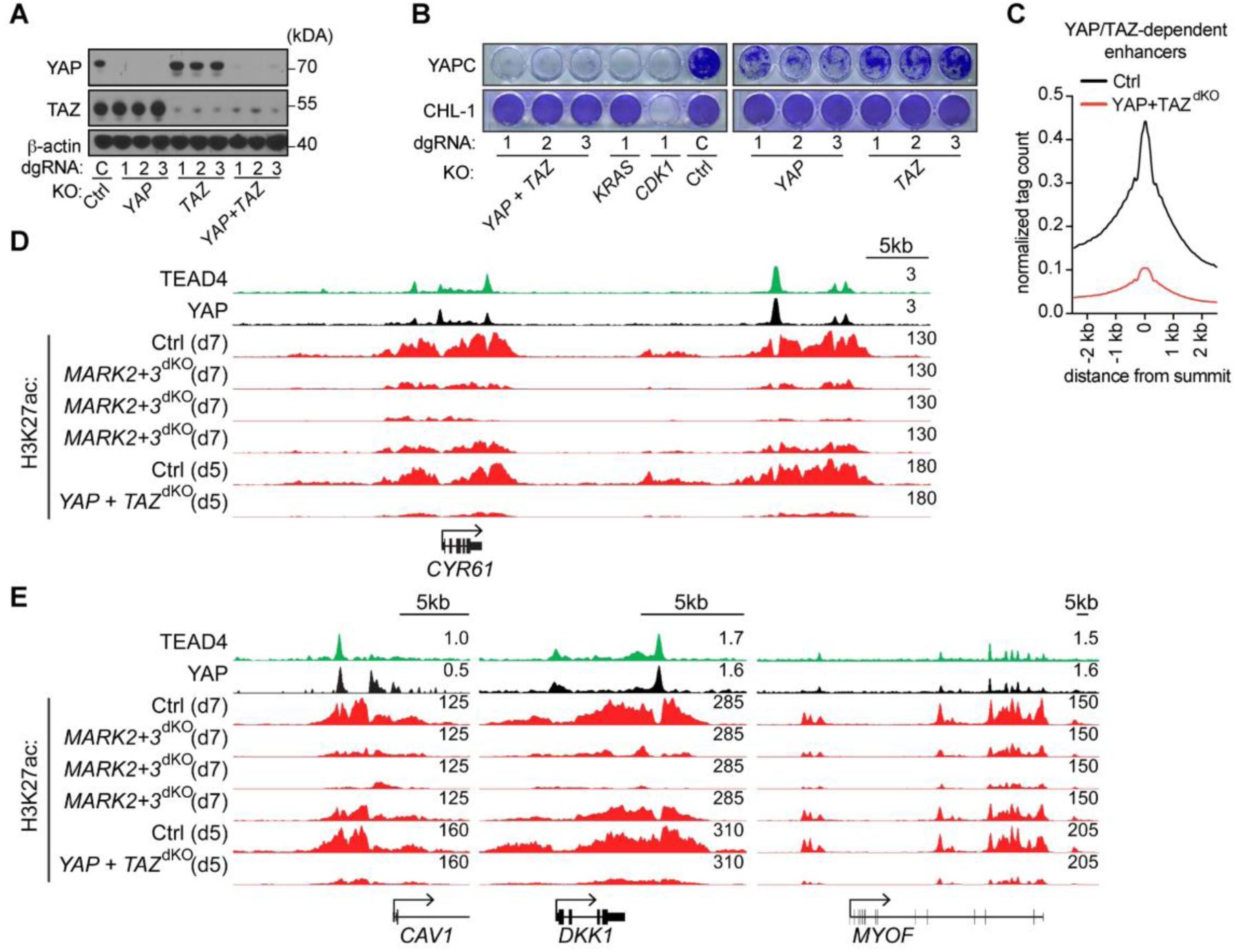
**A**, Western blot analysis in Cas9^+^ YAPC cells. **B**, Crystal violet stain of YAPC and CHL-1 (MARK2/3 independent) cells following dgRNA assisted lentiviral knockout of indicated genes. Data are representative of three independent experiments. **c**, CUT&RUN density profile of YAP/TAZ sensitive H3K27ac marked enhancer loci (n=7,896) following YAP+TAZ^dKO^. Profiles shown are an average of 50bp bins around the summit of the enhancers. **D**,**E** Occupancy profiles of public Chromatin immunoprecipitation sequencing (ChIP-seq) (TEAD4, YAP) (GSE66083) and CUT &RUN (H3K27ac) upon indicated gene knockout at YAP/TAZ target gene loci. (Three different dgRNAs for each *MARK2* and *MARK3)* (Data shown are an extension of Fig. 2I).

**Supplementary Fig. S3.**
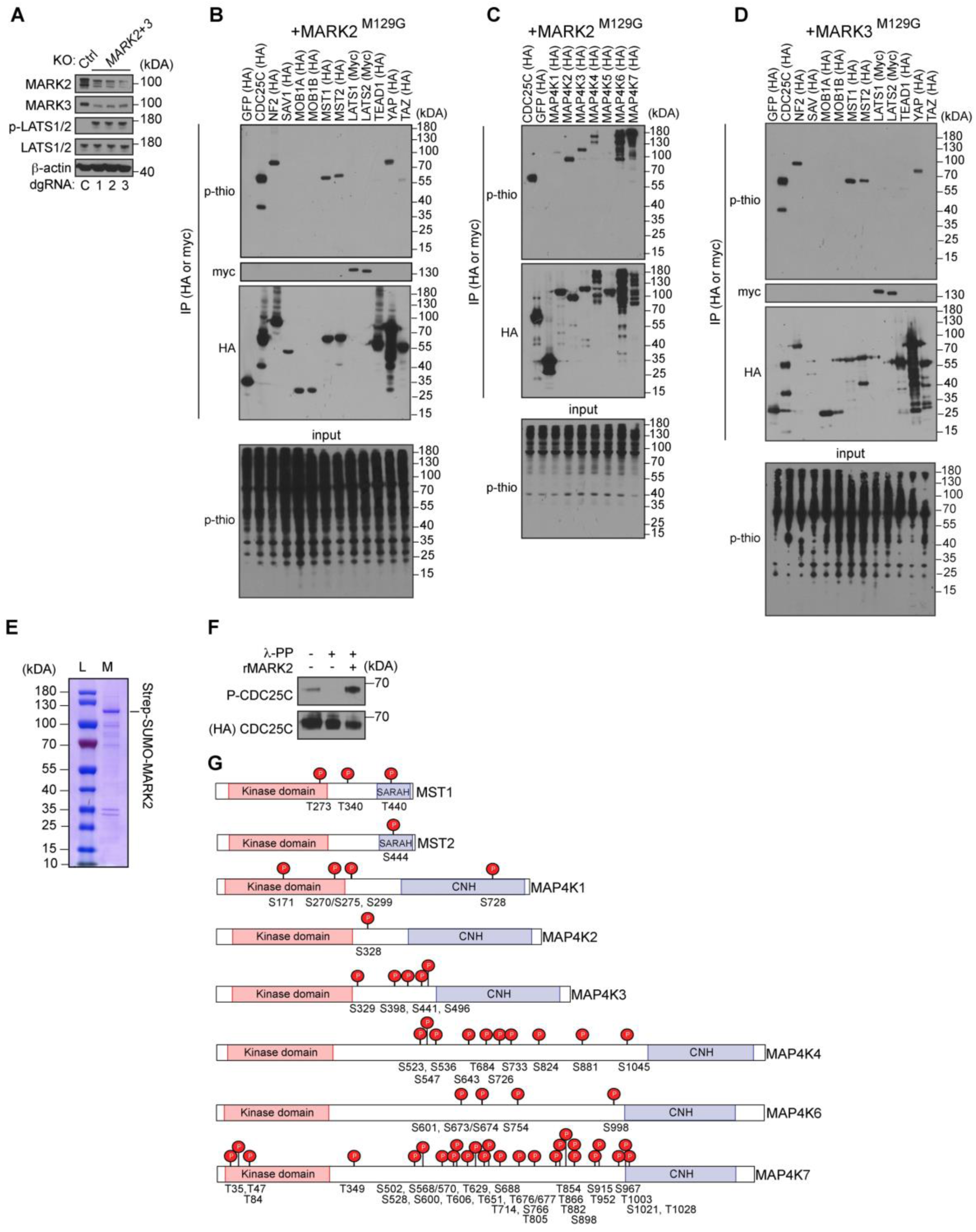
**A,** Western blot analysis of Cas9^+^ YAPC cells. **B**-**D**, Western blot analysis of b,c MARK2 or d, MARK3 specific in-cell phosphorylation of Hippo pathway components. Substrates were labeled as described in Fig. 3D, and phosphorylation was identified by staining with thiophosphate ester-specific antibodies. Data are representative of two independent experiments. **E**, Coomassie stain of affinity purified recombinant Strep-SUMO tagged MARK2 (rMARK2) purified from insect cells. **F**, Western blot analysis of purified HA-CDC25C following treatment of Lambda phosphatase (l-PP) and phosphorylation using rMARK2. **g,** Lolli-pop illustration of MARK2-dependent phosphorylation sites on MST1/2 and MAP4K1-4,6,7 identified using mass spectrometry-based phosphoproteomics. SARAH= Sav/Rassf/Hpo domain, CNH= Citron homology domain.

**Supplementary Fig. S4.**
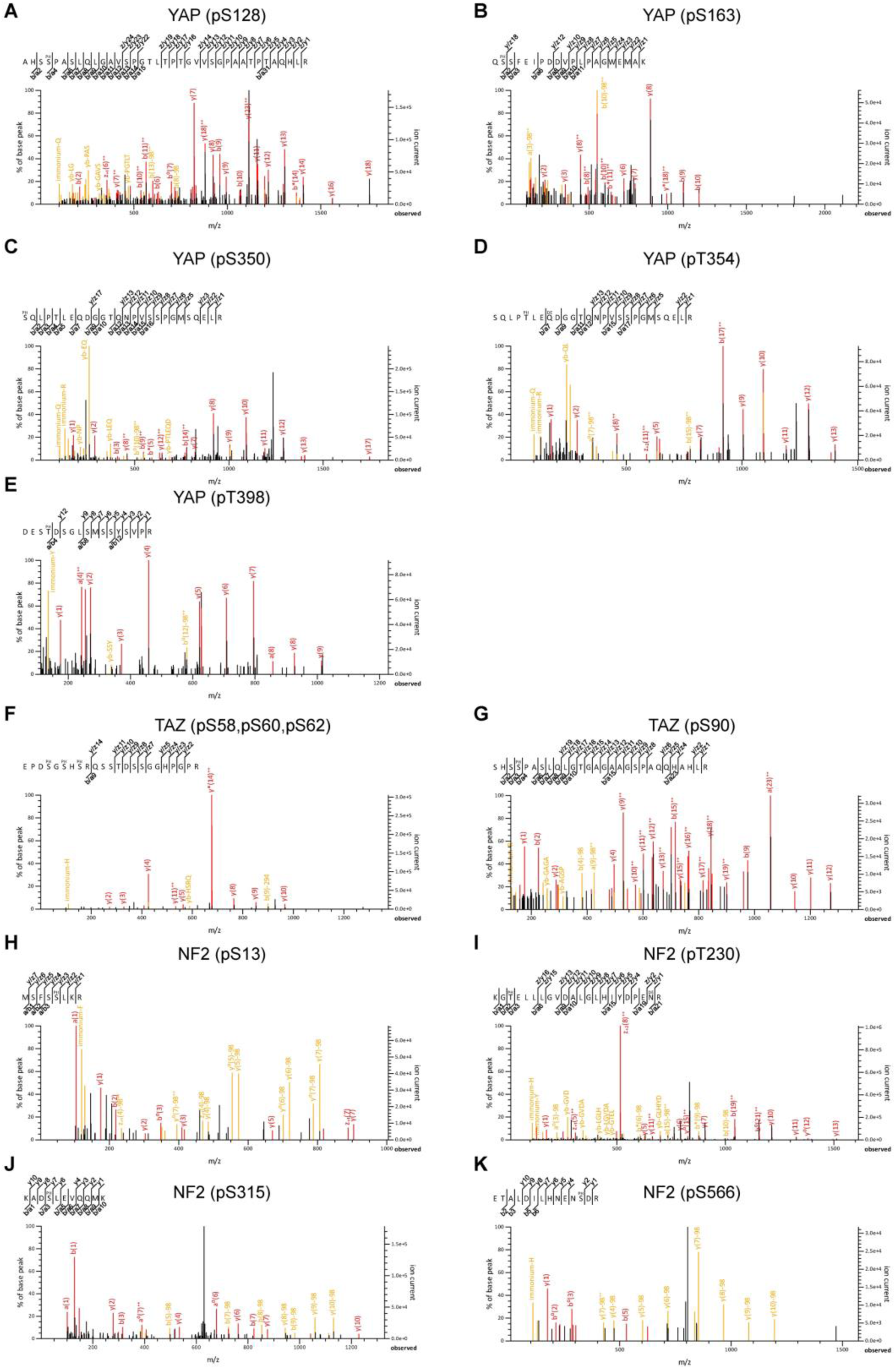
**A-K**, Fragmentation spectrum supporting peptide identity and phosphosites on YAP, TAZ, and NF2. Precursor ion chromatogram (XIC) and corresponding ions are provided in supplementary materials.

**Supplementary Fig. S5.**
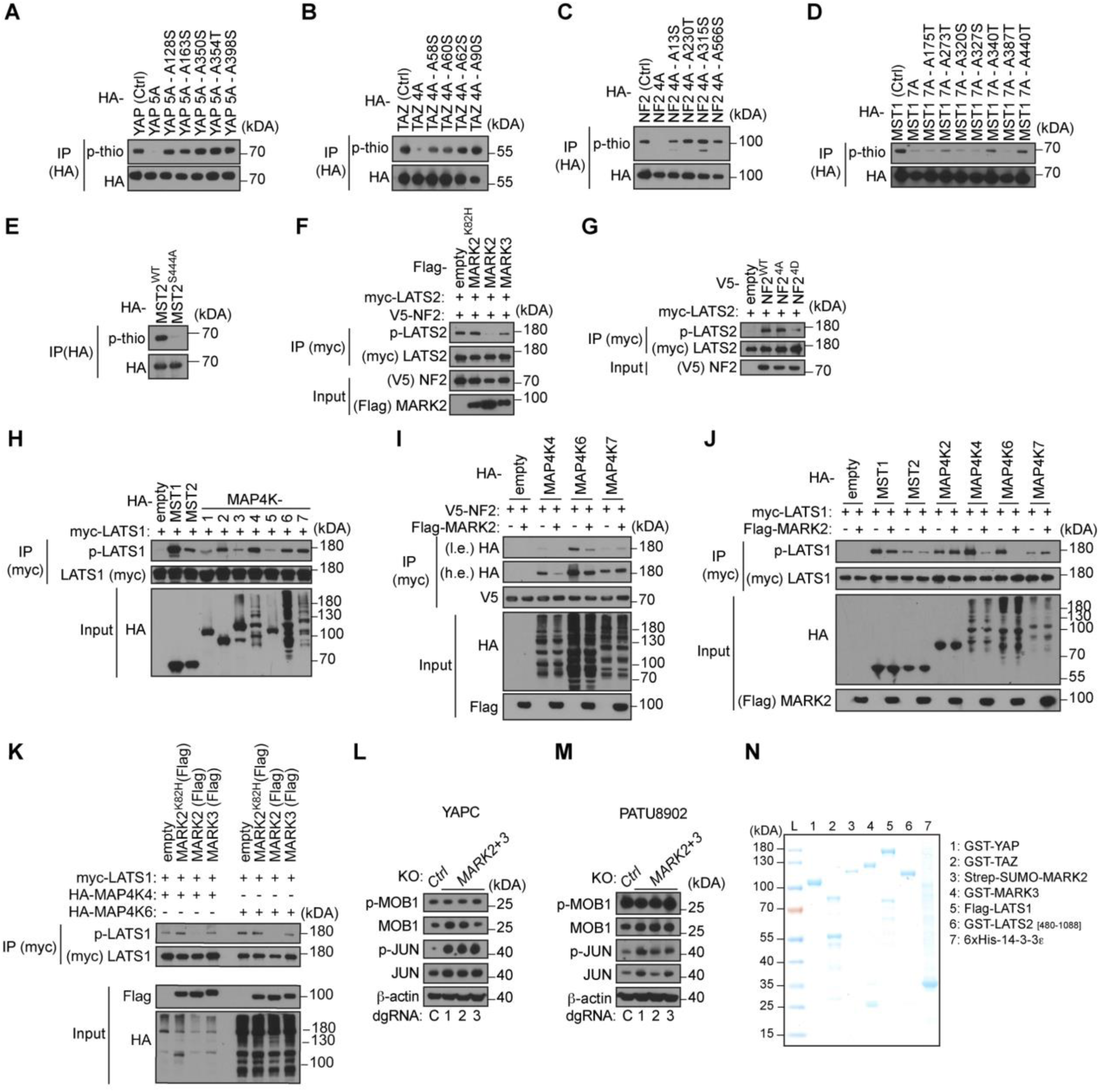
**A-E,** Western blot analysis of MARK2 specific substrates phosphorylation. Labeling as described in Fig 3d. Data are representative of two independent experiments. **F**, IP–western blot analysis evaluating the phosphorylation p-LATS2 (T1041) in presence or absence of MARK2 or MARK3 following NF2 overexpression in HEK-293T cells. Data are representative of two independent experiments. **G**, IP– western blot analysis evaluating the phosphorylation p-LATS2 (T1041) after NF2 mutant overexpression in HEK-293T cells. Data are representative of two independent experiments. **H,** IP–western blot analysis evaluating the phosphorylation p-LATS1 (T1079) following indicated gene overexpression in HEK-293T cells. **I**, IP–western blot analysis evaluating the interaction of NF2 and MAP4K4,6,7 in presence or absence of MARK2 overexpression in HEK-293T cells. Data are representative of two independent experiments. **J,** IP–western blot analysis evaluating the phosphorylation p-LATS1 (T1079) following indicated gene overexpression in HEK-293T cells. Data are representative of two independent experiments. **K,** IP–western blot analysis evaluating p-LATS1 (T1079) in presence of MAP4K4 or MAP4K6 together with MARK2,3, kinase dead MARK2^K82H^ or empty vector control overexpression in HEK-293T cells. Data are representative of two independent experiments. **L**,**M,** Western blot analysis of Cas9^+^ indicated cell lines. **N,** Coomassie stain of recombinant proteins used in *in vitro* kinase assays (Fig.3K**, 3L**), purified from bacteria (GST-YAP, GST-TAZ) and insect cells.

**Supplementary Fig. S6.**
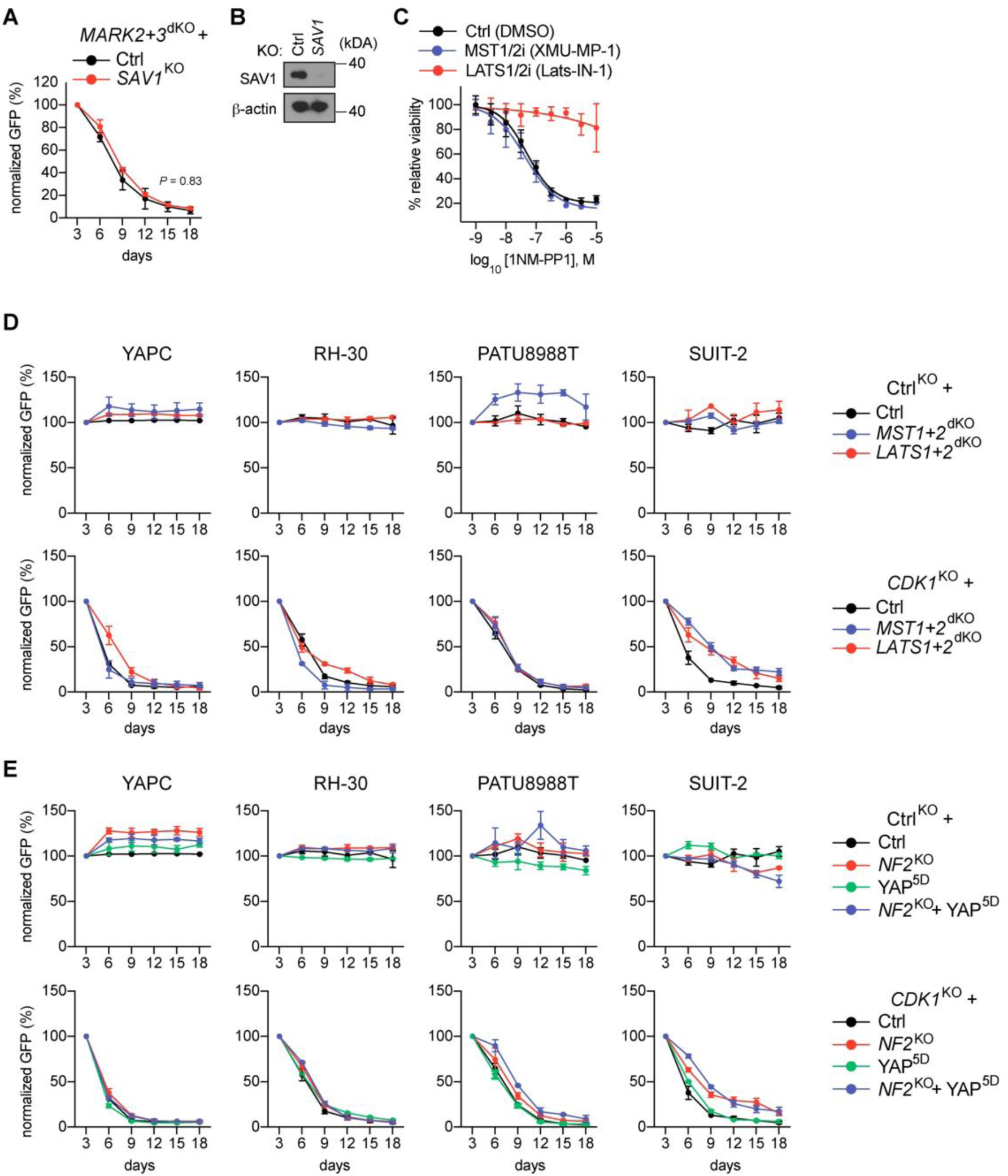
**A**, Rescue experiment of MARK2+3^dKO^ in Cas9^+^ YAPC cells following knockout of indicated genes. Data shown are the mean ± SD of %GFP^+^ (normalized to day 3 after infection). n=3-6. *P* values are calculated using a mixed effects model (considering the interaction of experimental groups over time) compared to Ctrl group and corrected with Bonferroni-Holm (BH). **B,** Western blot analysis of YAPC cells. **C**, Normalized relative luminescence units (RLU) from CellTiter-Glo viability measurements of Cas9^+^ YAPC-MARK2+3^dKO^ + MARK2^M129G^ cells following 5 days of combinational treatment of 1NM-PP1 and either +DMSO (0.1%), +500nM MST1/2 inhibitor (XMU-MP-1) or + 5µM LATS1/2 inhibitor (Lats-IN-1). Data are shown as mean ± SD. n=9 measurements from three biological replicates performed in triplicate. Four-parameter dose-response curves were plotted. **D**,**E** Competition-based fitness assays for Ctrl (dgRNA targeting hROSA26 locus) and knockout of essential gene *CDK1* corresponding to rescue experiment shown in Fig. 4A**, 4F**. Data shown are the mean ± SD of %GFP^+^ (normalized to day 3 after infection). n=3-6.

**Supplementary Fig. S7.**
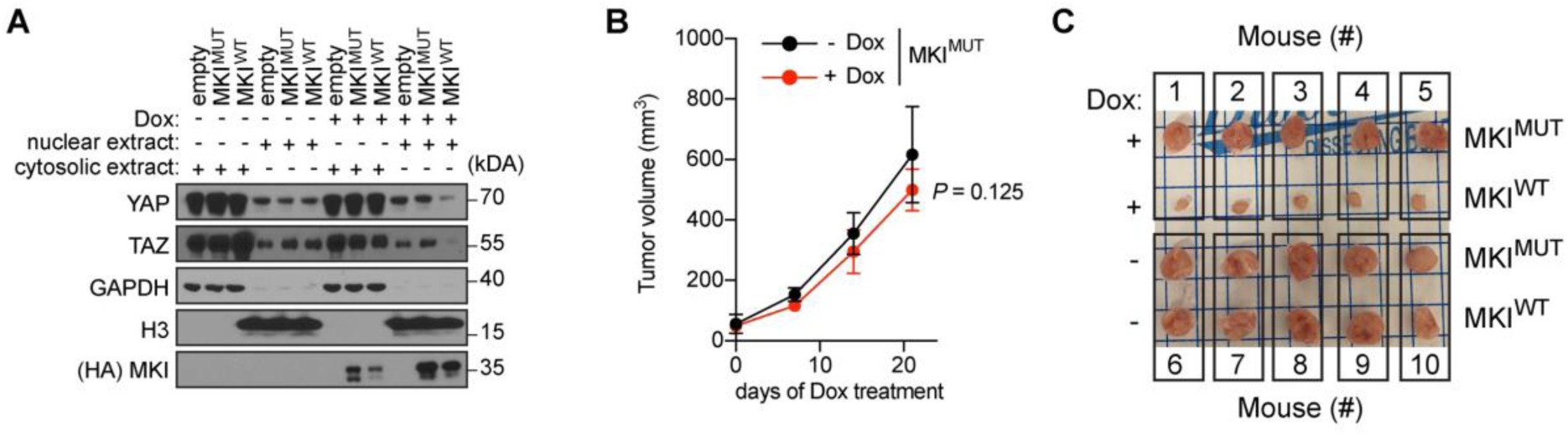
**A**, Western blot analysis of YAP localization following doxycycline (Dox) induced empty vector control, MKI^WT^ or MKI^MUT^ expression for 24h. Nuclear (Nuc) and cytosolic (Cyto) fractionation are indicated. (Data shown are an extension of Fig. 5E). **B,** Growth kinetics of subcutaneous YAPC xenografts implanted in immunodeficient mice. Expression of MKI^MUT^ from a Dox-inducible lentiviral construct was induced on day 10 post-injection of the cells. Data are shown as mean ± SD n=5 per group. *P* values are calculated using a mixed effects model (considering the interaction of experimental groups over time) compared to Ctrl group (-Dox) and corrected with Bonferroni-Holm (BH). **C**, Tumor imaging at the end-point of the xenograft experiments shown in **B**, and Fig. 5H.

## Contributions

O.K. and C.R.V. conceived this project and wrote the manuscript with input from all of the authors. O.K. and C.R.V. designed the experiments. O.K. performed experiments with help from D.S., C.T., A.A, F.M., D.A. and S.R.. O.K and T.H., performed statistical analysis. O.K. and O.E.D. designed CRISPR sgRNAs. O.K. designed and cloned paralog co-targeting CRISPR libraries. O.K. and D.S. screened libraries in cancer cell lines. O.K and C.T. performed experiments in subcutaneous xenografts. A.A. generated recombinant MARK2 proteins. F.M. and O.K. performed mass spectrometry sample preparations. F.M. and P.C. performed all mass spectrometry measurements. D.A., S.R. and O.K. performed organoid experiments. C.R.V., D.A.T., P.C., and D.L.S. supervised the studies and acquired funding.

